# MINDNet: Proximity interactome of the MICOS complex revealing a multifaceted network orchestrating mitochondrial biogenesis

**DOI:** 10.1101/2025.05.20.655052

**Authors:** Yulia Schaumkessel, Nahal Brocke-Ahmadinejad, Rebecca Strohm, Stefan Müller, Andreas S. Reichert, Arun Kumar Kondadi

## Abstract

The ‘Mitochondrial contact site and cristae organizing system’ (MICOS) complex is a multisubunit complex regulating mitochondrial inner membrane (IM) architecture, which is enriched at crista junctions (CJs) and required for cristae membrane dynamics. It modulates various mitochondrial processes including protein and lipid transport and is causally linked to a variety of human diseases. To gain a broad overview of the various pathways modulated by the MICOS complex, we examined its molecular neighbourhood. For this, we employed proximity biotinylation assays using APEX2 fused to four MICOS subunits (MIC10, MIC13, MIC26 and MIC27) in the respective mammalian knockout cells. These four MICOS-APEX2 fusion proteins integrated into the native MICOS complex and properly localised as revealed by electron microscopy combined with DAB staining and STED super-resolution nanoscopy. Here, we identify 119 common and 50 unique proteins, termed ‘**MI**COS **N**ano**D**omain **Net**work’ (MINDNet) encompassing the versatile proximity proteome of the MIC10/MIC13/MIC26/MIC27 subcomplex playing multifaceted mitochondrial functions. The MINDNet revealed a large number of OXPHOS proteins, protein translocases of the IM and OM, mitochondrial ribosomal proteins and solute carrier family transporters. Using the cues obtained from the proximity interaction studies, we investigated the role of all the MICOS proteins in modulating the function of the OXPHOS complexes. Among all the MICOS proteins, MIC10 and MIC60 consistently regulated the assembly and activity of the OXPHOS complexes. Overall, we propose that the MICOS complex integrates numerous spatial and temporal cues to regulate the dynamic microenvironment, along with IM architecture, which are involved in multiple pathways controlling mitochondrial biogenesis.

## Introduction

Mitochondria play critical roles in cellular homeostasis covering a broad range of functions such as energy conversion, inflammation, metabolic regulation, programmed cell death, epigenetic remodelling and nucleotide synthesis (Monzel *et al*, 2023; Suomalainen & Nunnari, 2024). The mitochondrial inner membrane (IM) is densely packed with proteins (Rath *et al*, 2021; Vögtle *et al*, 2017) and highly selective to various types of ions and metabolites entering the mitochondrial matrix. The IM is spatially demarcated into smooth inner boundary membrane (IBM), parallel to the outer membrane (OM), and cristae membrane (CM) invaginations by distinct physical entities present as pore-like structures around 25 nm in diameter termed crista junctions (CJs) (Daumke & van der Laan, 2025; Kondadi *et al*, 2020b; Zick *et al*, 2009). CJs act as diffusion barrier for membrane proteins resulting in compartmentalisation of proteins present in the IBM and CM (Gilkerson *et al*, 2003; Vogel *et al*, 2006; Wurm & Jakobs, 2006). While the CMs are enriched with the oxidative phosphorylation (OXPHOS) complexes, the IBM is enriched in proteins involved in mitochondrial import. It is proposed that CJs also regulate the movement of metabolites from the intracristal space (ICS, the lumen enclosed by individual cristae) to the remaining intermembrane space (IMS) and vice-versa (Frey *et al*, 2002; Mannella *et al*, 2001). In accordance, it was shown that CJs act as a barrier even for protons resulting in individual cristae with distinct levels of membrane potential (Wolf *et al*, 2019). CJs are enriched with proteins of the mitochondrial contact site and cristae organising system (MICOS) complex comprising two distinct subcomplexes namely MIC19/25/60 and MIC10/13/26/27 (Anand *et al*, 2016; Guarani *et al*, 2015). Deletion of MICOS proteins leads to loss or reduced formation of CJs separating the CMs and the IBM (Kondadi *et al*, 2020a; Stephan *et al*, 2020). Thus, the MICOS complex is a central component determining IM architecture (Kondadi *et al*., 2020b). Additionally, the MICOS complex was reported to play important roles in lipid trafficking, mitochondrial import, mitophagy and maintenance of mitochondrial DNA and membrane potential (Δψ_m_) (Kondadi & Reichert, 2024). Accordingly, it is not surprising that perturbations in mitochondrial IM structure are associated with a variety of pathologies including metabolic disorders and neurodegeneration (Daumke & van der Laan, 2025; Eramo *et al*, 2020; Kondadi & Reichert, 2024). Specifically, aberrations in MICOS complex subunits manifest as various pathologies. Mutations in *MIC10*, *MIC13*, *MIC26* and *MIC60* are associated with cerebellar atrophy, mitochondrial hepatopathy and mtDNA depletion syndrome (Kishita *et al*, 2024), fatal mitochondrial hepato-encephalopathy (Guarani *et al*, 2016), myopathy with cognitive impairment (Beninca *et al*, 2021), progeria-like phenotypes (Peifer-Weiss *et al*, 2023) and Parkinson’s disease (Tsai *et al*, 2018) respectively.

Although the MICOS complex is enriched at the CJs, the MICOS complex subunits interact with proteins possessing a tripartite localisation meaning the OM, the IM as well as the mitochondrial matrix. The following illustrations demonstrate the tripartite localisation of certain known MICOS interaction partners: 1) OM: MIC19 interacts with SAMM50 (a beta barrel protein) on one hand and MIC60 on the other hand (Darshi *et al*, 2011) forming the mitochondrial intermembrane space bridging (MIB) complex (Ott *et al*, 2012). MIC60 as well as MIC19 interact with solute carrier family 25 member 46 (SLC25A46), an OM protein. This interaction is required for maintaining mitochondrial lipid composition and cristae structure (Janer *et al*, 2016). 2) IM: MIC60 interacts with OPA1 (Glytsou *et al*, 2016) as well as MICU1 (Tomar *et al*, 2019) to regulate cristae morphology. Mic10 interacts with F_1_F_o_ ATP synthase to control respiratory growth apart from formation of CJs (Rampelt *et al*, 2022). Mic10 was also shown to interact with F_1_F_o_ ATP synthase subunit *e* Sue (Su *e*) in *Saccharomyces cerevisiae* (Eydt *et al*, 2017). 3) Mitochondrial matrix: MIC60 interacts with TFAM present in the nucleoids (Bogenhagen *et al*, 2008; Wang & Bogenhagen, 2006). Thus, despite indications that MICOS complex subunits could interact with different proteins localised to OM, various IM subcompartments, namely IBM and CM, as well as IMS, and matrix, studies shedding light on the molecular interactome of the MICOS complex are currently missing. In essence, the common as well as unique interactors of various subunits of the MICOS complex are poorly understood.

The classical affinity purification and mass spectrometry (MS) methods are not optimal for capturing weak or transient interactions and for discovering interaction partners mimicking the *in cellulo* or *in vivo* state. In order to overcome these limitations in our endeavour to decipher the interactome of the MICOS complex, we employed a proximity labelling assay using APEX2 (Ascorbate peroxidase 2). For this, HEK293 cells stably expressing *MIC10*, *MIC13*, *MIC26* or *MIC27*, and fused with APEX2 and Myc-tag on the C-terminus, were employed in the background of their respective knockout (KO) cells. Using Western blots (WBs), we found that HEK293 cells with stable exogenous expression of MIC10-, MIC13-, MIC26- or MIC27-APEX2-Myc (collectively termed MICOS-APEX2 from hereon) rescued the respective MICOS subunit levels. Further, the MICOS-APEX2 fusion proteins integrated into the MICOS complex as demonstrated using Blue-native (BN)-PAGE. 3,3’-diaminobenzidine (DAB) labelling of APEX2 fused to MICOS subunits revealed their localisation to IM using electron microscopy (EM). Similar to DAB staining, stimulated emission depletion (STED) super-resolution (SR) nanoscopy revealed the submitochondrial localisation of biotinylated proteins to the IM, IMS as well as the ICS. The interactome of the MICOS complex encompassed many proteins, localised to all mitochondrial subcompartments and performing a variety of mitochondrial functions. We found 119 common interactors of the MIC10/MIC13/MIC26/MIC27 subcomplex. 20 and 44 proteins were shared between any three or two MICOS subunits respectively. Interestingly, 50 unique interactors of the individual MICOS proteins were also found. Combined together, the sum of all the interaction partners comprises the ‘**MI**COS **N**ano**D**omain **Net**work’ (MINDNet in short) which could be broadly categorised into OXPHOS complexes, metabolite transporters, mitochondrial ribosomal proteins, regulators of protein import and quality control and mitochondrial dynamics demonstrating the diversity of the MICOS interactome. Probing into the functional interplay between MICOS and OXPHOS regulation, using a complete set of MICOS KO cell lines, we further demonstrated that MIC10 and MIC60 regulate the assembly and activity of the OXPHOS complexes. Thus, we propose that the MICOS complex acts as a molecular safeguard in regulating the assembly of OXPHOS complexes in addition to multiple other pathways required for mitochondrial biogenesis.

## Results

### Generation of four HEK293 cell lines stably expressing various MICOS-APEX2 fusion proteins

The MICOS complex forms two subcomplexes, the MIC10 and the MIC60 subcomplex, comprising MIC10/MIC13/MIC26/MIC27 and MIC19/MIC25/MIC60, respectively (Anand *et al*., 2016; Guarani *et al*., 2015). Here, we set out to understand the proximity interactome of the MIC10 subcomplex. For this, we made use of HEK293 cells deleted for *MIC10*, *MIC13*, *MIC26* or *MIC27*. These KO cell lines were used for stable expression of respective MICOS subunits fused to APEX2 followed by a Myc tag on the C-terminus namely MIC10-APEX2-Myc, MIC13-APEX2-Myc, MIC26-APEX2-Myc or MIC27-APEX2-Myc (collectively termed MICOS-APEX2 for simplicity from hereon) (**Fig 1A**). The amino acid sequence of linkers between various MICOS subunits and APEX2 are shown. Further, the predicted structures of MIC10, MIC13, MIC26 and MIC27 (blue colour) according to AlphaFold 3 protein structure database (Jumper *et al*, 2021) are depicted along with the APEX2 (red colour) and Myc tags (orange colour and shown using arrows) (**Fig 1A**). Since the proximity ambience of the MICOS proteins in the IM positions it in a unique spatial niche where they could interact with proteins present in the OM, IMS and IM on one hand and mitochondrial matrix proteins on the other hand including those destined for matrix, while passing through the mitochondrial IM and IMS, we reckon that using cells expressing matrix-targeted APEX2 and IM-targeted APEX2 as controls would lead to a robust investigation of the MICOS subcomplex interactome. Therefore, WT cells transiently expressing matrix-targeted APEX2, termed matrix-APEX2, were employed as done before (Sen *et al*, 2022) along with WT HEK293 cells stably expressing a transmembrane domain (TMD) targeted to the IM and fused to APEX2 and Myc tags on the C-terminus, termed IM-APEX2. The IM portion was generated using the N-terminal 116 amino acids of SCO1. For this, a section of MitoT plasmid was used (Viana *et al*, 2021). The predicted structures of matrix-APEX2 and IM-APEX2 are shown (**Fig 1A**). We standardised the conditions for performing APEX2-mediated proximity labelling in the extremely crowded IM using IM-APEX2 fusion protein (**Fig S1**). The proximity labelling of APEX2 depends on its catalytic activity defined by optimal amounts of biotin phenol (BP) substrate and hydrogen peroxide (H_2_O_2_). Therefore, we used cell lines stably expressing IM-APEX2 to test the profile of biotinylated proteins at different concentrations of BP as well as H_2_O_2_ (**Fig S1A and B**). We conclude that 0.5 mM of biotin-phenol (**Fig S1A**) as well as H_2_O_2_ (**Fig S1B**) were optimal for conducting our proximity labelling experiments. WBs using streptavidin revealed the pattern of the various MICOS biotinylomes. The biotinylation pattern of cells stably expressing four MICOS-APEX2 fusion proteins was remarkably similar indicating many common interactors while being different to matrix-APEX2 control as expected (**Fig S1C**). Noteworthy, the proximity biotinylome of IM-APEX2 matched the profiles of MICOS-APEX2 fusion proteins to a good extent indicating at least a partial match among their interaction profiles possibly due to high molecular crowding in the IM. A biotinylation scheme illustrating the MICOS-APEX2 fusion constructs along with the IM-APEX2 and matrix-APEX2 controls based on their mitochondrial localisation is depicted (**Fig 1B**). It is expected that while MICOS-APEX2 proteins are mainly enriched at the CJs, the artificial IM-APEX2 construct will randomly distribute within the IM.

**Figure 1.**
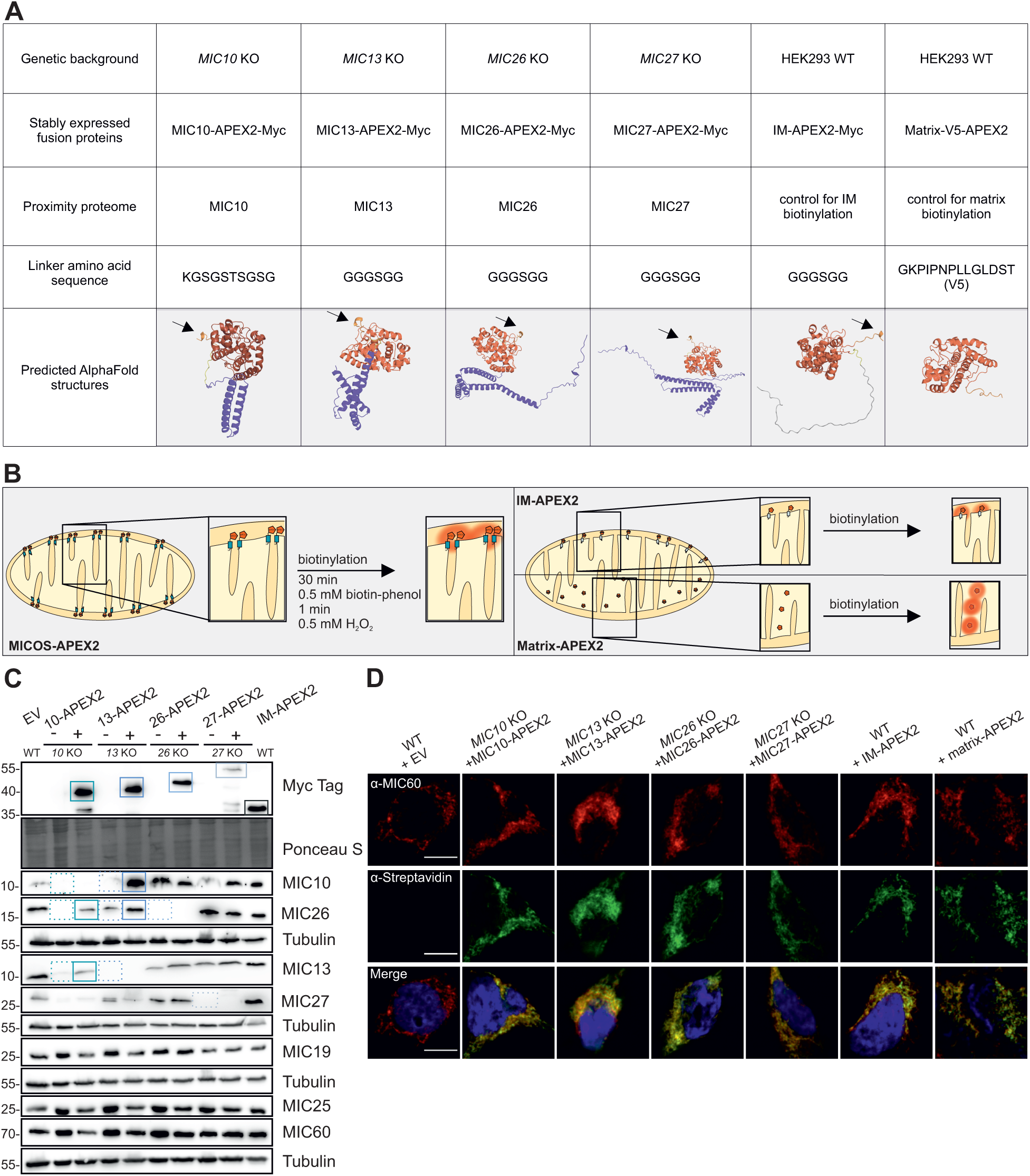
Generation of HEK293 cell lines stably expressing four different MICOS-APEX2 fusion proteins along with controls. (A) Schematic overview of various MICOS-APEX2-Myc fusion proteins (denoted as MICOS-APEX2 for simplicity from hereon) generated along with IM-APEX2-Myc (denoted as IM-APEX2 for simplicity from hereon) control fusion protein. Matrix-APEX2 was used as the second control condition. The amino acid linker sequences between respective MICOS proteins or IM transmembrane (TMD) domain and APEX2 are shown. AlphaFold 3 structure predictions, without cleavable mitochondrial targeting sequence, are represented using PyMol. Different MICOS subunits are depicted in blue colour. APEX2 and Myc tag are shown in red and orange (Myc tag additionally shown with arrows) respectively. The transmembrane domain of IM-APEX2 is depicted in light grey. The MICOS-APEX2 and IM-APEX2 fusion proteins were stably expressed in the background of respective MICOS KO and WT HEK293 cells, respectively. WT HEK293 cells, stably expressing empty vector (EV) were used as background control and, were employed for transient expression of matrix-APEX2. (B) Schematic representation of the APEX2 tagged to the respective MICOS subunit, IM-TMD and matrix controls. APEX2 catalyzes the biotin-phenoxyl radical production in the presence of biotin-phenol (BP) and H_2_O_2_ to initiate proteome tagging in the molecular neighborhood depicted in red rings. (C) Western blot (WB) analyses of various MICOS-APEX2 fusion proteins and IM-APEX2 control stably expressed in the corresponding MICOS KO background and WT cells, respectively. Stable expression of different MICOS-APEX2 fusion proteins results in restoration of the expected MICOS proteins (enclosed by solid rectangles) compared to corresponding KO cell lines (enclosed by dotted rectangles). All the MICOS proteins in cells expressing the respective MICOS-APEX2 along with Myc displayed increased molecular weight compared to the endogenous MICOS proteins as expected. Therefore, the rescue can only be seen using the anti-Myc antibody. Their expression is shown using the Myc antibody and the band at the expected molecular weight is additionally indicated with solid rectangles. (D) Fluorescence microscopy demonstrating the biotinylation activity of APEX2-constructs, using streptavidin conjugated to DyLight 488 within mitochondria marked using anti-MIC60 antibody in various cell lines mentioned. Nucleus is stained with Hoechst and shown in blue. Scale bar 10 µm.

First, we checked for the expression of various MICOS-APEX2 fusion proteins in the respective KO cell lines. WBs using an anti-Myc antibody confirmed the stable expression of all four MICOS-APEX2 fusion proteins (**Fig 1C**, enclosed using solid rectangles in topmost panel). It is known that deletion of *MIC10* (Kondadi *et al*., 2020a; Stephan *et al*., 2020) as well as *MIC13* (Anand *et al*., 2016; Guarani *et al*., 2015) leads to collective depletion of MIC10, MIC13 and MIC26 in multiple cell types. In congruence, our *MIC10 and MIC13* HEK293 KO cell lines show a loss of MIC10, MIC13 and MIC26 using the respective antibodies (**Fig 1C**, enclosed using dotted rectangles). Thus, stable expression of *MIC10* and *MIC13* should rescue the MIC10, MIC13 and MIC26 levels. In congruence, WBs using anti-MIC10 and anti-MIC13 antibodies reveal a rescue of the corresponding MICOS subunits upon stable expression of MIC10-APEX2 and MIC13-APEX2 in their respective KO cell lines (**Fig 1C**, enclosed using solid rectangles). Since, loss of neither *MIC26* nor *MIC27* affects other MICOS subunits, stable expression of MIC26 and MIC27 is expected to rescue only the corresponding proteins. Accordingly, the expression of MIC26-APEX2 and MIC27-APEX2 fusion proteins was confirmed in *MIC26* and *MIC27* KO cell lines using the anti-Myc antibody. Taken together, the stable expression of all four MICOS-APEX2 fusion proteins restored the amount of depleted MICOS subunits as expected indicating that the fusion proteins used are functional as they restored the level of those subunits normally depleted in the respective KO cells. Using the optimal standardised conditions for APEX2 labelling (**Fig S1**), we further checked for proper cellular targeting of MICOS-APEX2 fusion proteins (**Fig 1D**). For this, we determined the localisation of biotinylated proteins using streptavidin conjugated to DyLight 488. Fluorescence confocal microscopy revealed a clear colocalization of cells expressing MICOS-APEX2 to the mitochondria marked with an anti-MIC60 antibody. The IM-APEX2 and matrix-APEX2 controls were also correctly targeted to the mitochondria as the biotinylated proteins showed a mitochondrial localisation. Hence, we successfully generated a battery of four cell lines stably expressing MICOS proteins, fused to APEX2, in the corresponding KO background along with necessary control cell lines expressing IM-APEX2 or matrix-APEX2.

### MICOS-APEX2 fusion proteins localise to the mitochondrial inner membrane and integrate into native complexes

After confirmation of mitochondrial localisation of MICOS-APEX2 fusion proteins (**Fig 1D**), we further checked whether the generated MICOS-APEX2 fusion proteins expressed in the respective KO lines assemble into the native MICOS complex. The MICOS complex assembles into native complexes of different molecular weights (Huynen *et al*, 2016). Performing BN-PAGE electrophoresis on isolated mitochondria using antibodies against MIC10, MIC13, MIC26 and MIC27 revealed that the MICOS complex exists in different sizes in WT HEK293 cells (**Fig 2A-D**, lane 1) as observed previously (Naha *et al*, 2024). Deletion of *MIC10*, *MIC13*, *MIC26 and MIC27* led to no detection of the native complexes when probed with antibodies against respective proteins as expected (**Fig 2A-D**, enclosed by dotted rectangles). All MICOS-APEX2 fusion proteins successfully integrated into the native MICOS complexes restoring the endogenous pattern in the respective KO lines (enclosed using solid rectangles). Further, we used DAB staining to study the submitochondrial localisation of various MICOS-APEX2 fusion proteins in individual mitochondria. APEX2 has been routinely used to decipher spatial localisation by generating contrast while acquiring high-resolution EM images (Lam *et al*, 2015). In the presence of H_2_O_2_, APEX2 catalyses the oxidative polymerisation and local deposition of DAB after cellular fixation. EM images provide a sharp contrast due to recruitment of electron-dense osmium by the polymerised DAB. Control cells not expressing APEX2 displayed minimal electron-dense IM staining while the HEK293 cells stably expressing MIC10-, MIC13-, MIC26- and MIC27-APEX2 showed a clear positive staining pattern in the ICS as well as IMS suggesting that the MICOS fusion proteins possess a topological pattern as expected with the APEX2 present towards the IMS (**Fig 2E**). The IM-APEX2 control marked the ICS as well as the IMS while matrix-APEX2 control marked the matrix confirming proper localisation. In addition to the DAB staining, we performed STED SR imaging using antibodies against biotin and MIC60 marking the proximity proteome of MICOS-APEX2 fusion proteins and the MICOS complex respectively (**Fig 2F**). Consistent to the DAB staining, we also observed that the biotinylation pattern was prevalent at the edges of mitochondria (show using arrows) which was also colocalising with the MICOS complex. The IM-APEX2 biotinylome was spatially similar to the MICOS-APEX2 biotinylome and distinct from matrix-APEX2 control. Overall, using a set of validated experiments comprising a combination of SDS- and BN-PAGE, EM using DAB staining, confocal microscopy and STED SR nanoscopy, we confirm the successful generation of four different cell lines stably expressing four different functional MICOS-APEX2 fusion proteins along with IM-APEX2 control cell line.

**Figure 2.**
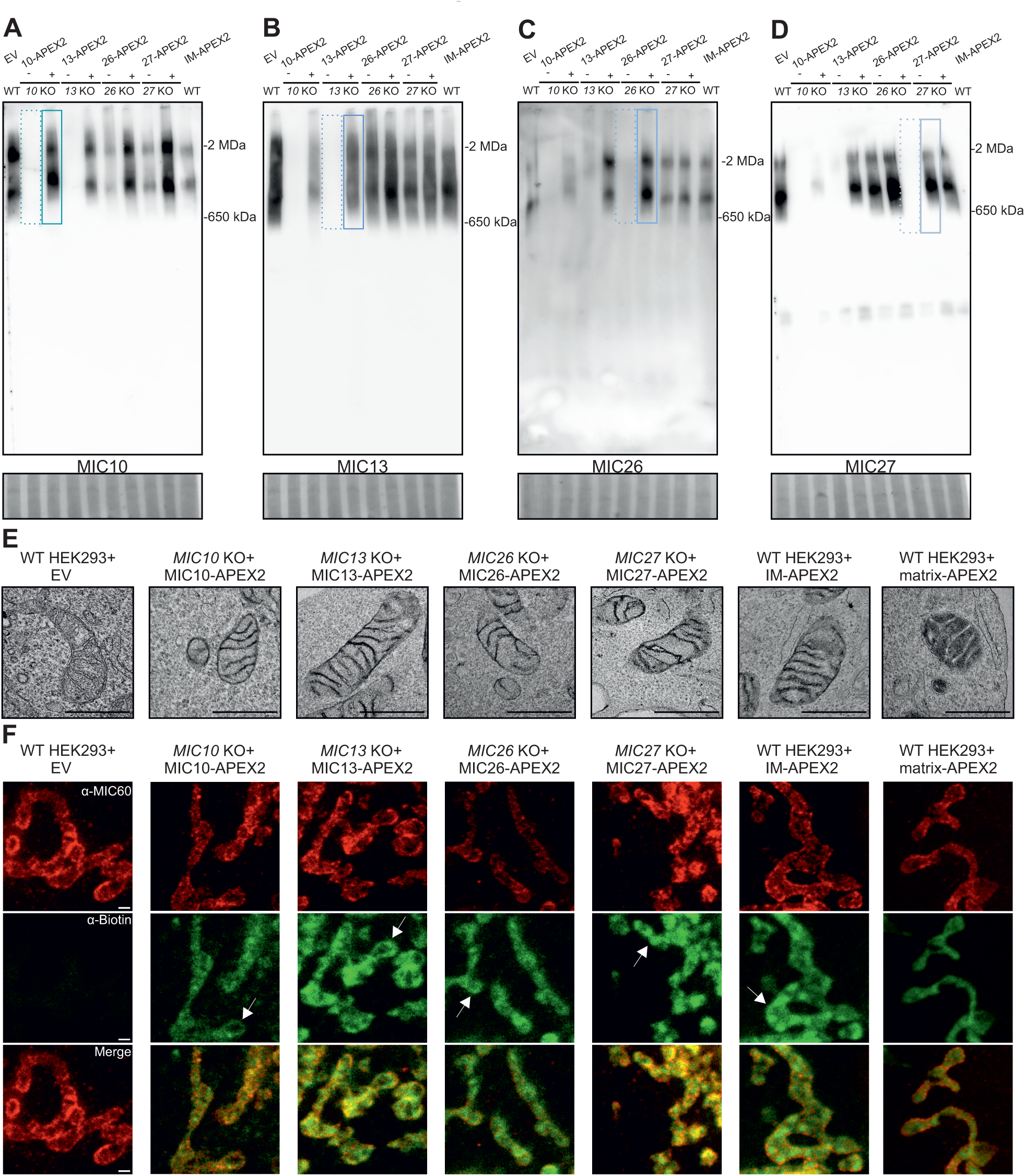
MICOS-APEX2 fusion proteins localise to the mitochondrial inner membrane and integrate into native complexes. (A – D) Blue Native (BN)-PAGE analyses of four cell lines stably expressing MICOS-APEX2 fusion proteins in the corresponding MICOS KO cell lines along with WT HEK293 cells expressing an empty vector (EV) or IM-APEX2 fusion protein used as control cell line. Efficient incorporation of (A) MIC10-, (B) MIC13-, (C) MIC26- and (D) MIC27-APEX2 fusion proteins (enclosed by solid rectangles) into the MICOS complex is observed using the respective antibodies. The dotted rectangles represent loss of various MICOS proteins in the respective KO cell lines. Coomassie stain is used as loading control comprising regions between 500 and 800 kDa. (E) Transmission electron microscopy (TEM) images demonstrating the MICOS-APEX2, IM-APEX2 and matrix-APEX2 submitochondrial localization in various cell lines. DAB oxidation catalysed by APEX2 results in local deposition of DAB polymer. All the MICOS-APEX2 fusion proteins as well as the IM control reveal the DAB staining in the intermembrane space (IMS) as well as intracristal space confirming the orientation of APEX2 towards the IMS. The matrix-APEX2 reveals matrix DAB staining as expected. Scale bar 500 nm. (F) Images acquired by STED super-resolution nanoscopy demonstrate the pattern of biotinylation in individual mitochondria resulting from MICOS-APEX2 fusion proteins. Biotinylation is prevalent at the rim of mitochondria as indicated by white arrows. The IM-APEX2 reveals similar biotinylation pattern to MICOS-APEX2 different to matrix-APEX2. The MICOS complex, enriched at the CJs, is marked using an anti-MIC60 antibody. Scale bar 500 nm.

### Analysis pipeline and MICOS interactome organized according to their submitochondrial localization and mitochondrial functions

For deciphering the interactome of one of the two MICOS subcomplexes, we employed cell lines stably expressing MIC10-, MIC13-, MIC26- or MIC27-APEX2 fusion proteins in the respective KO background, along with control cell lines expressing either IM-APEX2 or matrix-APEX2, using five replicates for each condition. Treatment of the above-mentioned six cell lines with BP and H_2_O_2_ was followed by quenching and cell lysis. All the cell lysates were used to coimmunoprecipitate (coIP) the biotinylated proteins with streptavidin as described before (Tan *et al*, 2020). The captured biotinylated proteins were subjected to label-free quantitative proteomics approach. Principal component analysis (PCA) plots are commonly used to analyse the variation among and between different groups in large proteomics datasets. We observed that various groups of MICOS-APEX2 fusion vectors were grouped similarly while the matrix-APEX2 control assembled as a separate group distant from the MICOS-APEX2 proteomics dataset (**Fig S2A**). The IM-APEX2 control although grouped distinctly was still similar to MICOS-APEX2 fusion proteins, possibly due to the IM crowding, indicating the IM localisation of IM-APEX2 and MICOS-APEX2 proteins. (**Fig S2B**). A scheme representing the workflow of our enrichment studies by proximity labelling is depicted (**Fig 3A).** The MS yielded two classes of enriched interaction partners: First, those which were detected in a sufficient number of technical replicates (≥3 out of 5) and enriched in the biotinylome of the respective MICOS-APEX2 group when compared to the matrix-APEX2 or IM-APEX2 controls, mentioned as enriched (log_2_FoldChange (FC) category) from now on. Second, those which were exclusively found in the biotinylome of the respective MICOS-APEX2 but not in in the matrix-APEX2 or IM-APEX2 controls, defined as enriched (**F**ound **O**nly in **M**ICOS-APEX2, **FOM** category in short) from now on. Regarding this, we observed that the six APEX2 cell lines including the controls subjected to proteomics had a similar distribution of number of replicates (out of five replicates) where the proteins were detected ruling out any possible bias in the analysis of enriched proteins in the FOM category for matrix-APEX2 and IM-APEX2 for the whole cell (**Fig S2C**) and mitochondrial proteins (**Fig S2D**).

**Figure 3.**
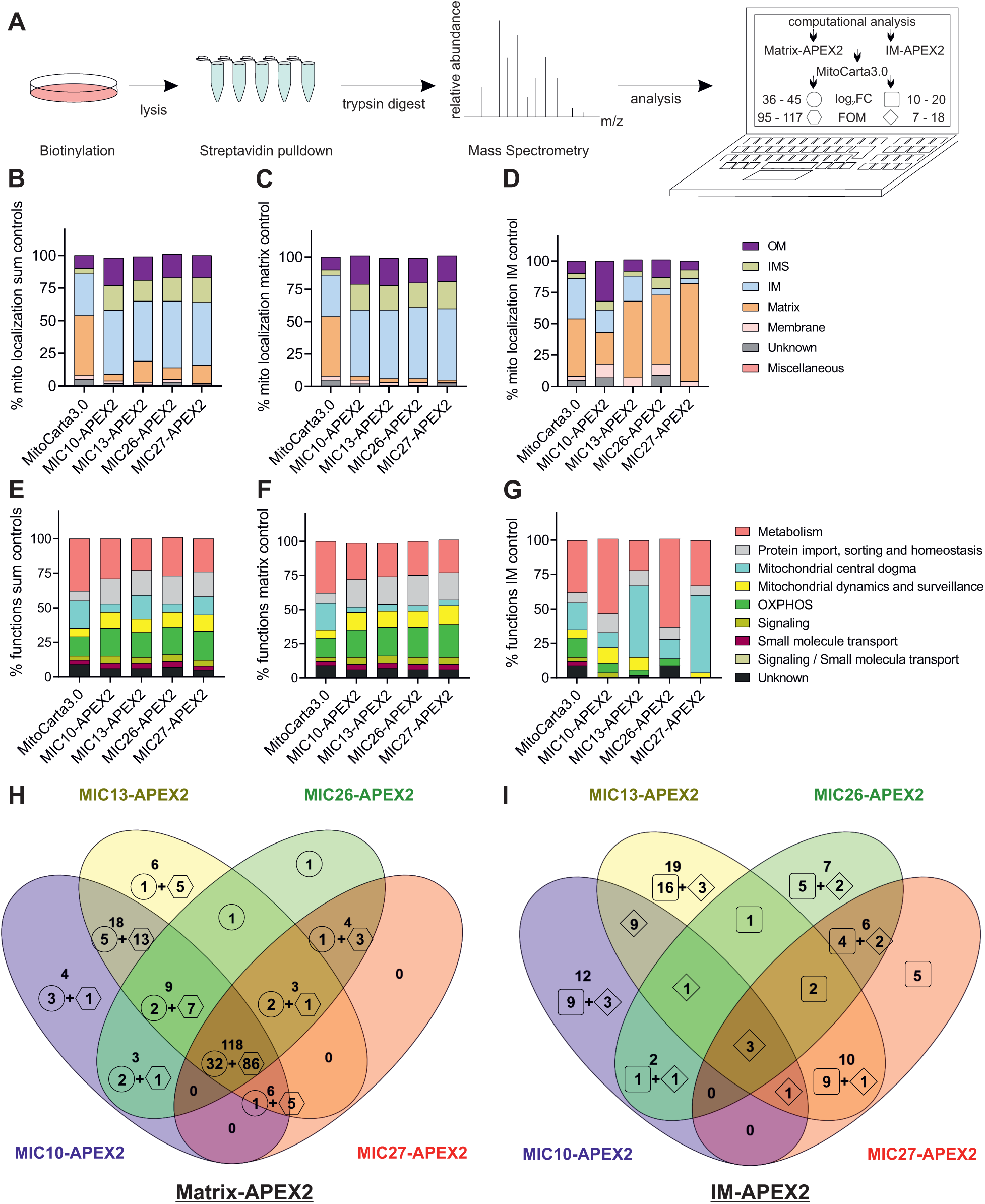
Analysis pipeline and MICOS interactome organized according to their submitochondrial localization and mitochondrial functions. (A) Schematic representation of the experimental setup and the analysis pipeline. The MICOS interactors were obtained after normalization over the matrix-APEX2 and IM-APEX2 controls separately upon MitoCarta3.0 filtering. 95 to 117 MICOS interactors (FOM (**F**ound **O**nly in respective **M**ICOS) proteins absent in the control, represented using a hexagon) and 36 to 45 MICOS interactors (log_2_FC enriched proteins over control condition, represented using circle) were uncovered upon normalization over matrix-APEX2 control whereas 7 to 18 (FOM category, represented using square diamond) and 10 to 20 MICOS interactors (log_2_FC enrichment, represented using square) were uncovered upon using IM-APEX2 as control. (B – D) Histograms depicting the percentage of proteins possessing the respective submitochondrial localization assigned according to MitoCarta3.0 compared to the interactome of various MICOS subunits. Histograms representing the mitochondrial localization of the total sum of interactors obtained upon combining interactomes normalized to both controls (B), matrix-APEX2 control (C) and IM-APEX2 control (D). Annotated mitochondrial subcompartments include outer membrane (OM), intermembrane space (IMS), inner membrane (IM), matrix, membrane, unknown and miscellaneous compartment and are shown using different colour-codes. (E – G) Histograms depicting the percentage of proteins possessing the respective mitochondrial functions assigned according to MitoCarta3.0 compared to the interactome of various MICOS subunits. Histograms representing the mitochondrial functions of the sum total of interactors obtained upon combining interactomes normalized to both controls (E), matrix-APEX2 control (F) and IM-APEX2 control (G). Various mitochondrial functions are shown using different colour-codes and comprise the following categories: metabolism, protein import, central dogma, dynamics and surveillance, OXPHOS, Signaling, small molecule transport and unknown. (H and I) Venn diagram depicting the number of common enriched proteins among the MIC10/MIC13/MIC26/MIC27 interactome in different combinations obtained using matrix-APEX2 (H) and IM-APEX2 (I) as controls, respectively. The category of FOM proteins (**F**ound **O**nly in **M**ICOS and absent in the control) is represented using a hexagon and square diamond obtained upon using matrix-APEX2 and IM-APEX2 controls, respectively. Log_2_FC enriched proteins over control, are represented using circle and square upon using matrix-APEX2 and IM-APEX2 control, respectively.

Overall, we detected 4,245 proteins across all the datasets combined (**Table S1**). 4,119 proteins remained after removing reverse and potential contaminants. Out of the total of 4,119 proteins for all conditions, 2,613 to 2,892 proteins were detected in the MIC10-, MIC13-, MIC26- and MIC27-APEX2 conditions (**Fig S3A**, step 1). From these, a total of 1,181, 1,606, 1,146 and 1,250 proteins were significantly enriched (*p* value ≤ 0.05, log_2_FC ≥ 0 and FOM categories together) in the respective conditions mentioned above (**Fig S3B and C**). Since the mitochondrial localisation and catalytic activity of the generated APEX2 fusion proteins was confirmed using electron and confocal microscopy as well as STED SR nanoscopy (**Fig 1D, 2E and F**), we focussed on the mitochondrial proteins according to the list from MitoCarta3.0 (Rath *et al*., 2021) using matrix-APEX2 (**Table S2**) and IM-APEX2 as controls (**Table S3**). From this, we obtained 176 to 239 enriched mitochondrial proteins in our dataset comprising all the four MICOS-APEX2 conditions **(Fig S3A, step 2)**. It must be kept in mind that MICOS-APEX2 interactors enriched over the respective controls (matrix-APEX2 or IM-APEX2) were already distributed into two categories, enriched log_2_FC (proteins present in respective controls) and exclusively found FOM (proteins absent in respective controls) category. Thus, the enriched proteins found only in MICOS-APEX2 and absent in respective controls (FOM group) were not influenced by the log_2_FC cut-offs set for the two controls (**Fig S3A**, step 2 to step 3). Regarding the matrix-APEX2 control, we used a standard cut-off of log_2_fold increase ≥1 for defining biologically meaningful enriched proteins when normalising the interactions of MICOS-APEX2. Regarding the IM-APEX2 control, we had observed using STED SR nanoscopy that the pattern of biotinylation obtained was remarkably similar to MICOS-APEX2 conditions (**Fig 2F**) along with staining pattern of biotinylated proteins detected using WBs (**Fig S1B and C**). This and the reasoning that the mitochondrial IM is a densely proteinaceous structure allowed us to relax the cut-off to a log_2_fold enrichment of ≥0.5 **(Fig S3A)**. After using the respective cut-offs and combining the data from both the control samples, a total of 151 to 192 proteins were significantly enriched in the four MICOS-APEX2 conditions (*p* value ≤ 0.05 and log_2_fold increase ≥ 0.5 and 1 for IM-APEX2 and matrix-APEX2 controls respectively, including the FOM category proteins). These 151 to 192 mitochondrial proteins comprised a significant fraction of 16.3 to 22.3 % of the total captured interactome when all cellular proteins were taken into account (log_2_fold enrichment of ≥ 0.5 or 1 for IM- and matrix-APEX2 control respectively, and the FOM category) (**Fig S3D**).

The overall uniformity of the four MICOS interactomes used in this study is evident upon distribution of the interactors based on the submitochondrial localisation and different mitochondrial pathway functions (**Fig 3B and E**) defined according to MitoCarta3.0 (Rath *et al*., 2021). We rationalised that an increase in percentage of proteins in the MICOS interactome compared to MitoCarta3.0 upon considering the submitochondrial localisation and mitochondrial pathway functions indicates selective enrichment of the respective proteins in the MICOS neighbourhood. Considering the submitochondrial localisation of interactome of all four MICOS subunits upon filtering with MitoCarta3.0, we found that there was a robust increase in percentage of proteins localised to IM, IMS and OM while there was decrease in percentage of MICOS interactors localising to mitochondrial matrix (**Fig 3B**). Considering the known localisation of the MICOS proteins, this observation underpins the general validity of the approach used. The percentage of proteins localising to different submitochondrial compartments when using matrix-APEX2 (**Fig 3C**) and IM-APEX2 (**Fig 3D**) as controls was also analysed separately showing an expected shift depending on the control used. Next, we analysed the datasets with respect to various functional categories provided by the Mitocarta3.0 database. Here, we observed a clear enrichment in the percentage of proteins playing vital roles in OXPHOS, protein import, sorting and homeostasis and mitochondrial dynamics and surveillance, when compared to the percentage of these proteins to MitoCarta3.0 (**Fig 3E**). Again, the percentage of proteins performing different mitochondrial functions is shown separately when using matrix-APEX2 (**Fig 3F**) and IM-APEX2 (**Fig 3G**) as controls. Venn diagrams demonstrate the number of common interactors among MIC10-, MIC13-, MIC26-as well as MIC27-APEX2 obtained using matrix-APEX2 (**Fig 3H**) as well as IM-APEX2 (**Fig 3I**) controls. We observe a large number of proteins (118 interactors) present as common interactors while using matrix-APEX2 as the normalising control (**Fig 3H**). On the contrary, when compared to the IM-APEX2 control, the number of common interactors for all four MICOS subunits, namely 3, was much lower (**Fig 3I**) pointing to a more subunit-specific local protein neighbourhood. Although the mitochondrial IM is enriched with MICOS proteins at the CJs, it does not exclude the presence of MICOS proteins in the rest of the IM i.e., CM and the IBM. Therefore, we reasoned that using matrix-APEX2 as a control allows us to find the majority of the common interactors of the MICOS complex whereas using IM-APEX2 as control may specifically rather uncover MICOS interactors at the nanodomains of CJs to reveal unique interactors of each of the MICOS proteins examined.

### The common MICOS interactome includes OXPHOS, transporters and import machinery proteins and engages with proteins involved in a broad range of molecular functions

The MitoCarta3.0 provides an annotation of all the 1136 proteins by classification into 7 different pathways, namely, metabolism (427), central dogma (230), OXPHOS (160), protein import, sorting and surveillance (83), mitochondrial dynamics and surveillance (70), small molecule transport (34 genes), signalling (31) and other unknown functions (101) (Rath *et al*., 2021). Our classification of the MICOS interactors (using different colour-codes) is strictly based on their assigned function in MitoCarta3.0. Thus, the total of 119 common interactors for all four MICOS subunits, namely MIC10, MIC13, MIC26 and MIC27 result from the combination of comparisons for both controls, since two out of three common proteins found using IM-APEX2 as control were also found in the 118 common proteins using matrix-APEX2 control. These 119 proteins comprising the MINDNet are further shown using a graphical depiction of mitochondrial membranes by sorting into seven different categories based on their functions using different colour-codes (**Fig 4**). Out of the 119 common interactors of the MICOS subcomplex, the largest category of proteins belonged to OXPHOS and metabolism followed by protein import, sorting, homeostasis category containing 28, 28 and 22 proteins, respectively. These three categories comprised 65.5 % of the total common interactors. The majority of the OXPHOS proteins being common MICOS interactors belonged to complex IV (18 proteins) followed by complex I and III with 7 and 3 common interactors, respectively. The next largest group of common MICOS interactors (28 proteins) belonged to the proteins involved in metabolism and included transporters of the solute carrier family (SLC)25A (8 proteins) including SLC25A12 and SLC25A13 (**Fig 4**). These two proteins along with glycerol-3-phosphate dehydrogenase 2, mitochondrial (GPD2), are part of the malate-asparate shuttle and the glycerol-3-phosphate shuttle, respectively, which play important roles in NADH/NAD^+^ redox balancing between the cytosol and the mitochondrial matrix suggesting an important metabolic function of the MICOS complex. Other interactors in this category include sideroflexins (SFXNs) 1 and 3, oxygen-dependent coproporphyrinogen-III oxidase, mitochondrial (CPOX) and StAR-related lipid transfer domain protein 7 (STARD7) playing vital roles in mitochondrial serine transport (Kory *et al*, 2018), heme biosynthesis pathway (Yien & Perfetto, 2022), coenzyme Q transport, and ferroptosis (Deshwal *et al*, 2023). Further, the protein import and homeostasis (including quality control pathways) category contained 22 common MICOS interactors such as translocases of outer membrane (TOMM) 40 and 70, translocases of inner membrane (TIMM) 9, 10, 13, 17B, 23 and 50 as well as IM proteases OMA1 and YME1L. In addition, 16 common MICOS interactors belong to the category of proteins involved in mitochondrial dynamics and surveillance pathways include syntaxin 17 (STX17), DIABLO and FKBP Prolyl Isomerase 8 (FKBP8). All the MICOS proteins were present emphasizing the interaction of MICOS proteins within the complex validating our study. Some other notable interactors of the MICOS complex with those involved in signaling pathway comprise voltage dependent anion channel (VDAC) 1, 3, mitochondrial antiviral-signaling protein (MAVS) and mitochondrial calcium uptake (MICU) 1 and 2 and AAA Domain Containing 3 (ATAD3) A and B. Altogether, the above-mentioned classes of proteins comprise the common interactome of the MICOS complex demonstrating a diverse and dynamic microenvironment of the mitochondrial IM necessary to meet the ever-changing metabolic cellular demands.

**Figure 4.**
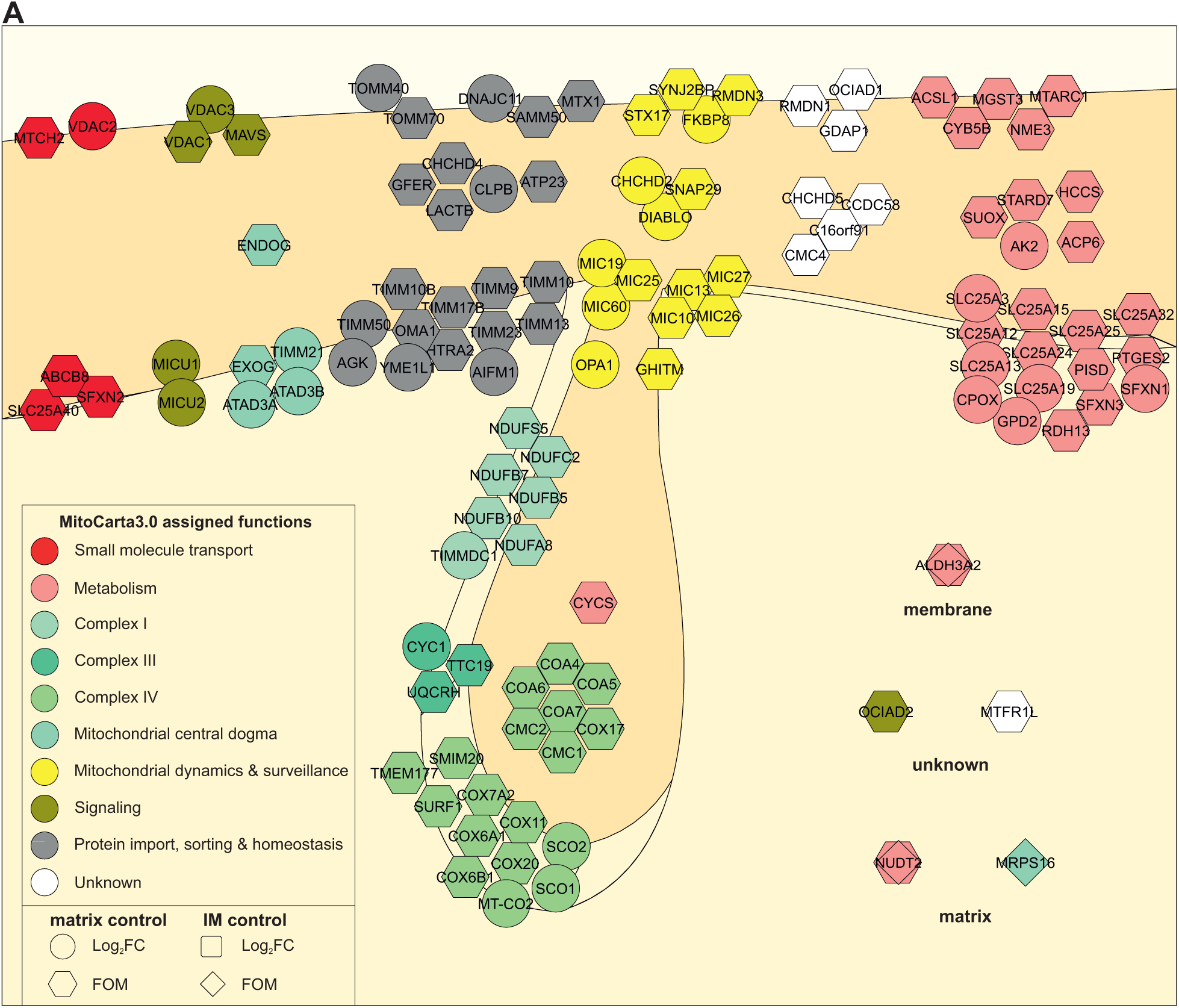
The versatile ‘<u>MI</u>COS <u>N</u>ano<u>D</u>omain <u>Net</u>work’ (MINDNet) engages with proteins involved in a broad range of molecular functions including OXPHOS, solute carrier (SLC) family transporters and mitochondrial import machinery proteins. Schematic depiction of the common interactors of MICOS-APEX2 fusion proteins. The different interactors were colour-coded and grouped according to the functions assigned in MitoCarta3.0 and placed in the respective mitochondrial subcompartment comprising OM, IMS, IM, matrix, membrane and unknown compartment. The shape of different nodes consisting of the respective gene names was also kept consistent to the previous figure. Different MICOS subunits (preys) were present in the mitochondrial dynamics and surveillance category (yellow colour) as common interactors of all four MICOS-APEX2 fusion proteins as expected. The MINDNet includes 119 common candidates. A sum of 118 and three common candidates were obtained using matrix-APEX2 and IM-APEX2 controls, respectively. Two of the candidates were enriched in all MICOS-APEX2 in comparison to both controls (indicated by superimposed shapes) taking the total tally to 119 common proteins.

### Overlapping and unique interactors of MIC10, MIC13, MIC26 and MIC27

Our analysis reveals that 68.2 % of proteins were found as common interactors of the MIC10/MIC13/MIC26/MIC27 subcomplex upon using matrix-APEX2 control while 55.1 % of the proteins (majority) maintained unique interactions with individual MICOS proteins upon comparison to IM-APEX2 control (**Fig 5A**). Thus, the IM-APEX2 control acts as specific spatial nanodomain control while the matrix-APEX2 acts more as a general control. 20 proteins were present in the interactome of any three of the MICOS-APEX2 fusion proteins and sorted according to MitoCarta3.0 based on their functions and colour-codes maintained from the previous figure (**Fig 5B**). The interactors of the respective MICOS proteins are present within the nodes and connected by lines. 18 interactors were obtained using matrix-APEX2 as control and 4 proteins were obtained using IM-APEX2 as control, with two proteins overlapping between both controls, namely CASP8 and FKBP10. The interactions in this section revealed some well-studied proteins like prohibitins and myosin 19 (MYO19). Proteins involved in protein import, sorting and homeostasis according to MitoCarta3.0 classification represent the largest subgroup in this category. Further, a total of 44 proteins (26 and 28 were obtained using matrix-APEX2 and IM-APEX2 controls respectively, with 10 proteins overlapping between both controls) were shared between any two of the MICOS-APEX2 fusion proteins among MIC10-, MIC13-, MIC26- and MIC27-APEX2 (**Fig 5C**). Here, the majority of the proteins belonged to the category ‘metabolism’ and included transporters like SLC25A11, SLC25A21 and SLC25A33. Mitochondrial ribosomal proteins like MRPL35, MRPL48, MRPL55 and MRPS22 were also part of the MICOS-APEX2 interactome. Lastly, we focussed on the proteins which were exclusively interacting with only one of the four MICOS-APEX2 fusion proteins (**Fig 5D**). We found that MIC10, MIC13, MIC26 and MIC27 interact uniquely with 14, 23, 8 and 5 proteins respectively. Some unique interactors of MIC10 include TOMM70, Ras Homolog Family Member T1 (RHOT1 or MIRO1), Mitochondrial Fission factor (MFF), Dynamin 1 Like (DNM1L or DRP1) and Peroxiredoxin 4 (PRDX4). The majority of the unique interactors of the MIC13 include mitochondrial ribosomal proteins like MRPL3, MRPL11, MRPL14, MRPL15, MRPL16 and MRPS9 and transporters SLC25A4 or ANT1, SLC25A20 exchanging ADP/ATP and acyl-carnitines across the mitochondrial IM respectively. Acetyl-CoA Carboxylase Alpha (ACACA), Aldehyde Dehydrogenase 1 Family Member L1 (ALDH1L1) and ALDH1L2 are some unique interactors of MIC26 while Histidyl-tRNA Synthetase 2, mitochondrial (HARS2), Catalase (CAT), MRPL46, MRPS25 and Tu Translation Elongation Factor, mitochondrial (TUFM) are unique interactors of MIC27. The unique interactors of different MICOS proteins belonging to the same MICOS subcomplex indicate their non-redundant functions.

**Figure 5.**
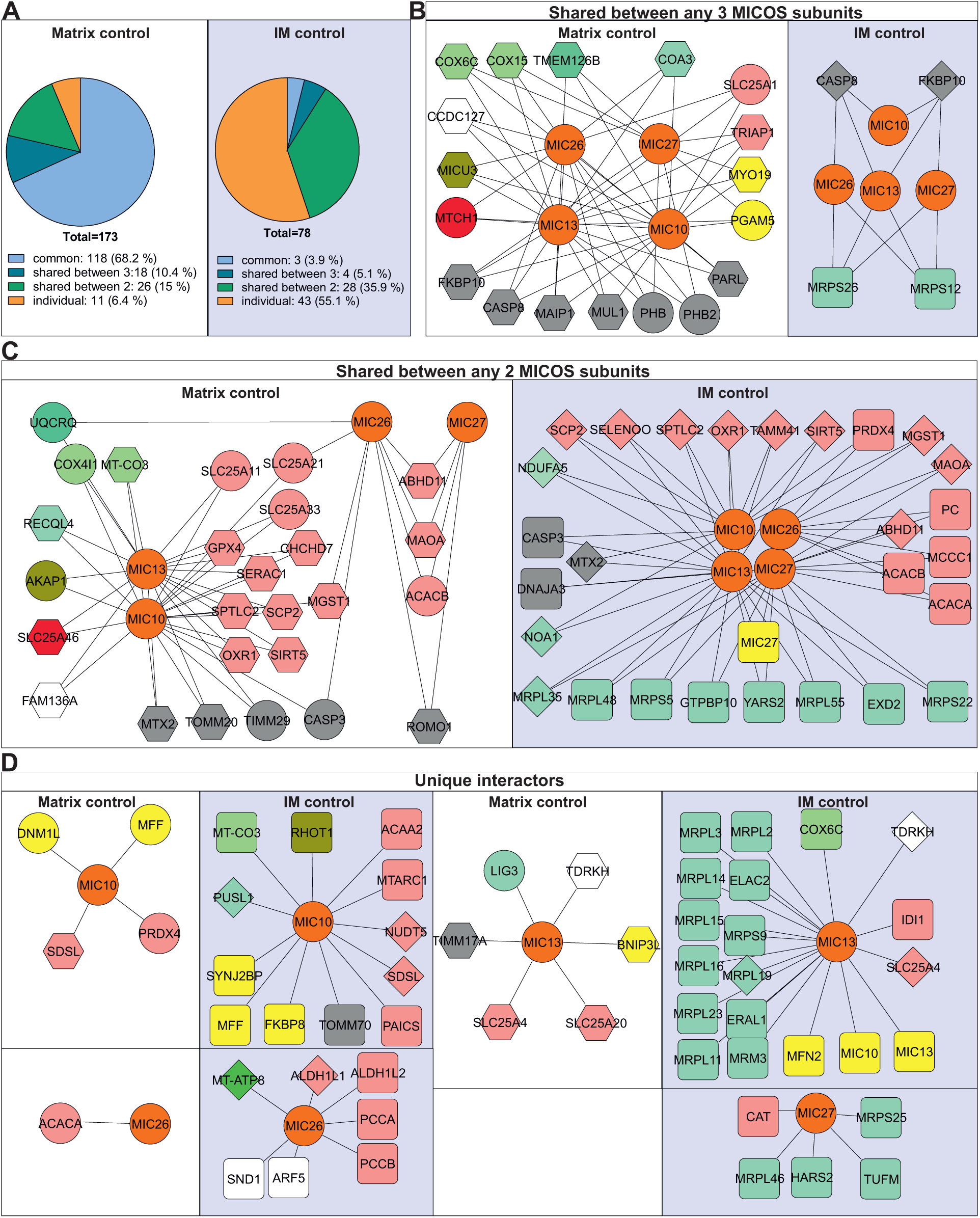
Overlapping and unique interactors of MIC10, MIC13, MIC26 and MIC27. (A) Pie charts representing the percentage of enriched hits among the various combinations of MICOS-APEX2 groups. The majority of enriched proteins (68.2 %) are common for all MICOS-APEX2 fusion proteins obtained upon using matrix-APEX2 control. Conversely, a majority of the enriched proteins are unique interactors (55.1 %) for MIC10-, MIC13-, MIC26- and MIC27-APEX2 fusion proteins when compared to the IM-APEX2 control. (B and C) Proximity proteome representation of enriched hits shared between any three (B) or any two (C) MICOS-APEX2 groups upon using the respective APEX2 control. (D) Unique interactors of the respective MICOS-APEX2 fusion proteins upon using the respective APEX2 control are shown. The proteins were colour-coded in accordance to the functions assigned in MitoCarta3.0 and the node shapes represent the affiliation to Log_2_FC enriched or FOM category as in previous figures. The interactions are shown by lines connecting the different nodes where the gene names of the interactors are used. MICOS subunits present in yellow represent the preys, whereas MICOS subunits in orange represent the employed APEX2-fused baits.

The complete list of enriched proteins with log_2_fold increase ≥ 1 (where the respective proteins were present in the control conditions) obtained using matrix-APEX2 (**Fig 6A**) as well as the enriched candidates with log_2_fold increase ≥ 0.5 when using IM-APEX2 (**Fig 6B**) control are shown using colour-coded intensity-based heat maps. Interestingly, OPA1 was present as the topmost interactor of all the 4 MICOS proteins investigated in this study when matrix-APEX2 was used as the normalisation control (**Fig 6A**). Additionally, volcano plots of all mitochondrial enriched interactors of the respective four MICOS interactomes belonging to this category are shown for matrix-APEX2 control (**Fig S4A-D**) and IM-APEX2 control (**Fig S4E-H**). A majority of the interactors obtained using IM-APEX2 were unique (**Fig 6B**) and not common among the 4 MICOS interactomes. Accordingly, the use of IM-APEX2 normalisation control resulted in clear distinction of colour-coded intensity-based map of log_2_fold increase when compared to matrix-APEX2 (**Fig 6A**). Further, the log_2_fold change candidates with relevant values and the enriched proteins belonging to the FOM category (where the respective proteins were absent in the control conditions) are shown in **Table S4**.

**Figure 6.**
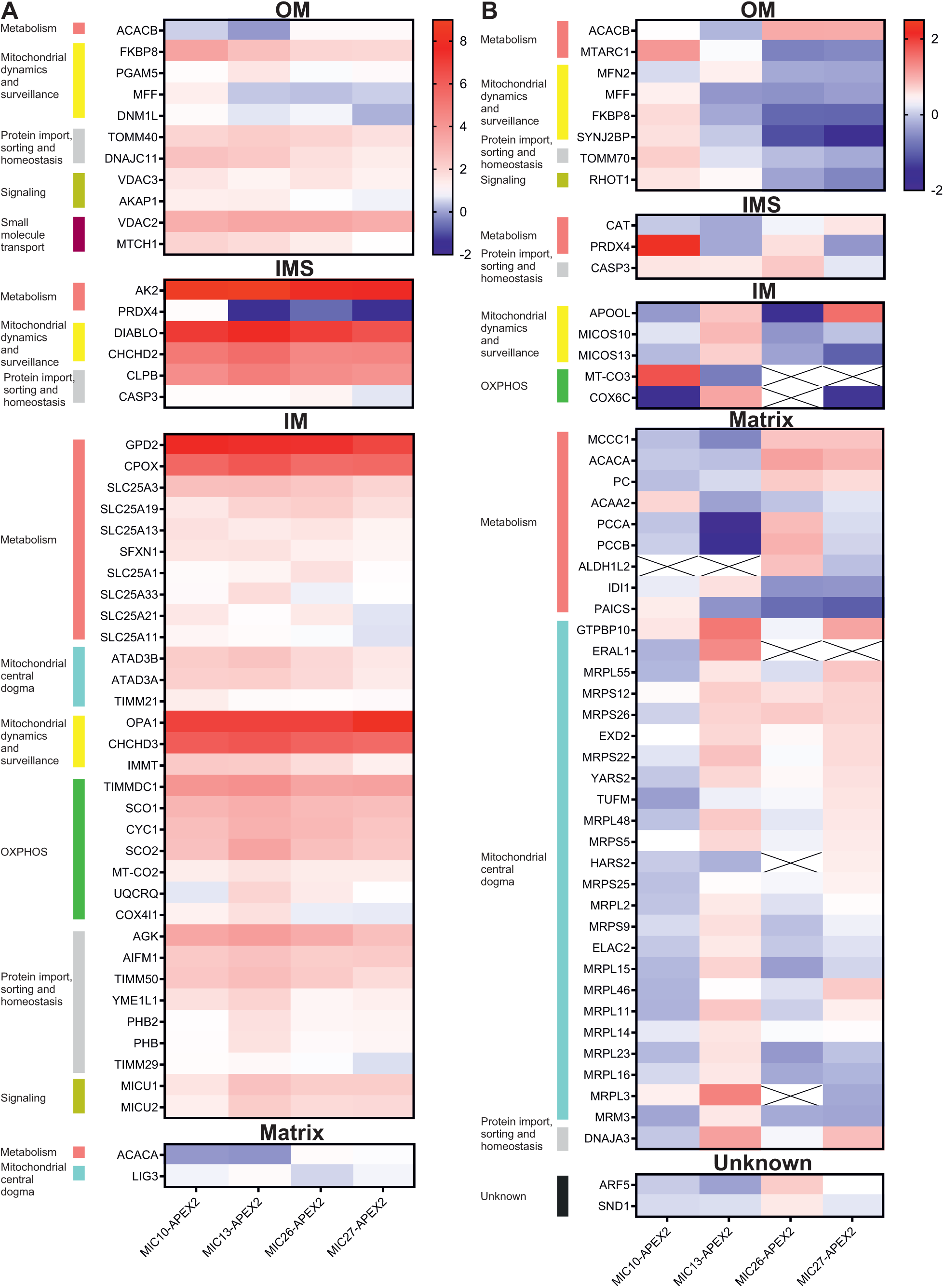
Heatmap showing the various MICOS interactors obtained using log_2_fold enrichment (excluding FOM category) (A and B) Heatmap representing the MIC10-, MIC13-, MIC26- and MIC27-APEX2 interactomes obtained using matrix-APEX2 (A) and IM-APEX2 (B) as controls. The heatmaps of the corresponding interactomes are distributed into mitochondrial subcompartments and functions. Missing proteins are crossed out.

### MIC10 subcomplex maintains the assembly and activity of OXPHOS complexes I and IV

The common interactors of the MICOS proteins used in this study revealed that the largest class of proteins (28 proteins) belonging to the OXPHOS complexes with complex I and IV comprising 7 and 18 proteins respectively (**Fig 4**). Therefore, we were encouraged to investigate the mutual regulation between OXPHOS assembly, at both individual complex and supercomplex (SC) level, and MICOS subunits. Utilising all the MICOS KO cell lines, WBs performed using antibodies against different MICOS proteins (**Fig S5A**) along with a detailed quantification (**Fig S5B**) reveal loss of specific proteins in respective MICOS KO cell lines in accordance with previous literature (Naha *et al*., 2024; Stephan *et al*., 2020). Then, we checked the assembly of the native OXPHOS complexes in all MICOS KOs using antibodies directed against proteins belonging to OXPHOS complexes I (NDUFB4), III (UQCRC2), IV (COXIV) and V (ATP5A) (**Fig 7A-D**). We found that the assembly of complex I was reduced in *MIC10*, *MIC13* and *MIC60* KO cells. Complex IV assembly was reduced in *MIC10* and *MIC60* KO cells whereas complex III and V (monomer) assembly was largely unchanged. Thus, MIC10 and MIC60 loss led to reduced assembly of complex I and IV while MIC13 loss specifically reduced the complex I assembly (enclosed by dotted rectangles).

**Figure 7.**
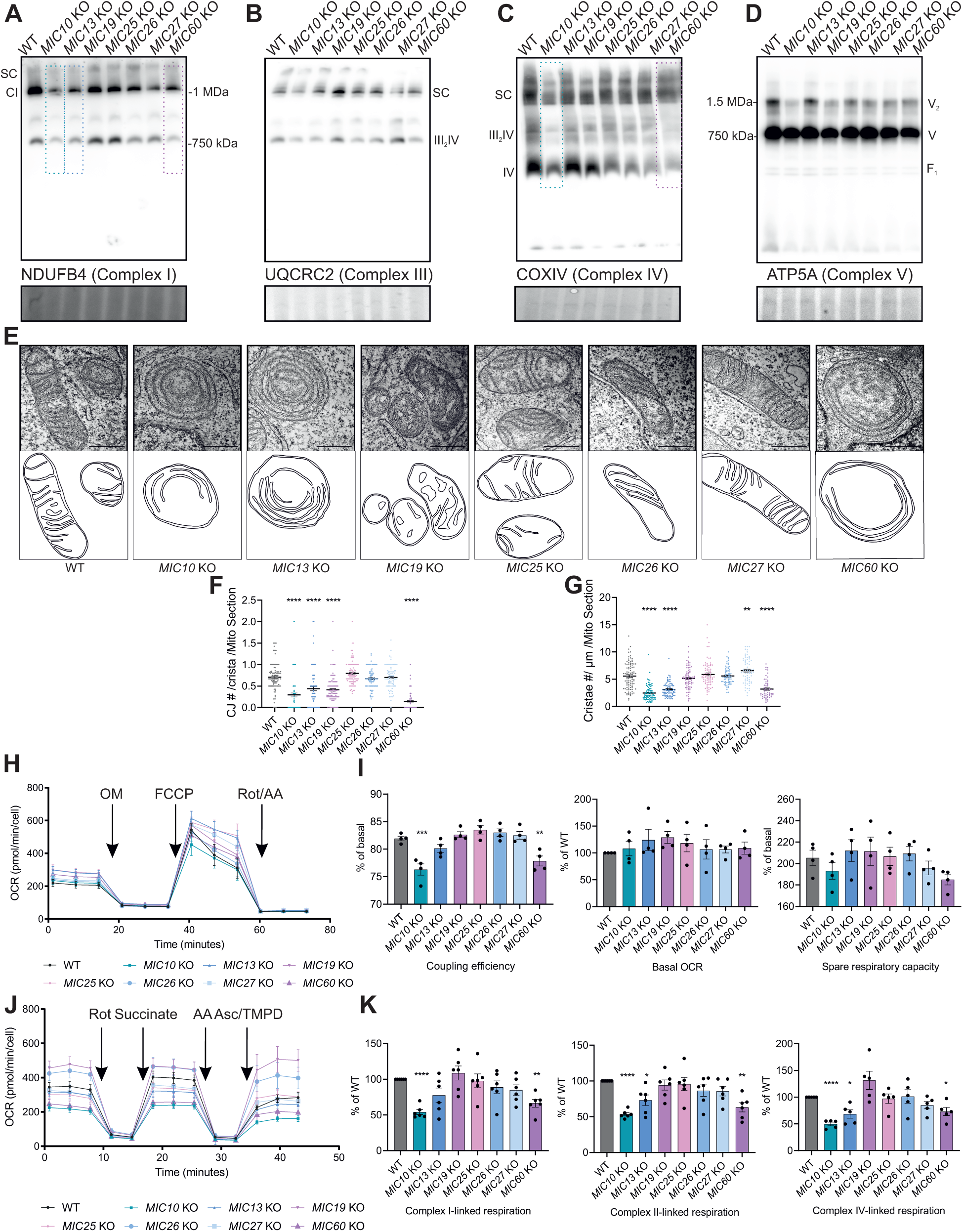
MIC10 subcomplex maintains the assembly and activity of OXPHOS complexes I and IV. (A – D) BN-PAGE analyses reveals a consistent reduction of OXPHOS complex I and IV assembly in *MIC10* and *MIC60* KO cells among all MICOS KO cells (indicated by dotted rectangles). Coomassie stain is used as loading control comprising regions between 500 and 800 kDa. (E – G) Representative TEM images demonstrating the mitochondrial ultrastructure of individual MICOS KO cells (E). Quantification reveals a significant reduction in the number of crista junctions (CJs) per crista in *MIC10* KO, *MIC13* KO, *MIC19* KO and *MIC60* KO cells (F) and cristae number per unit length (µm) in *MIC10* KO, *MIC13* KO and *MIC60* KO cells (G), (n = 36-79) (N = 2). (H and I) Representative mitochondrial stress test assessed with Seahorse XF analyzer using sequential injection of oligomycin, FCCP and rotenone/antimycin a (n = 8-9) (H). Quantification from various biological replicates shows a significant decrease in mitochondrial coupling efficiency in *MIC10* KO and *MIC60* KO cells (N = 4) (I). (J and K) Representative individual complex feeding run as assessed in permeabilized cells with Seahorse XF analyzer by sequential injections of rotenone, succinate, antimycin a and ascorbate/TMPD (n = 8-9) (J). Quantification of different biological replicates reveals a significant decrease of complex II and complex IV activities in *MIC10* KO, *MIC13* KO and *MIC60* KO, whereas complex I activity is significantly decreased only in *MIC10* KO and *MIC60* KO cells (K) (N = 6). Data are represented as mean ± SEM (H – K). Statistical analysis was performed using one sample *t*-test with \**P* < 0.05, \*\**P* < 0.01, \*\*\**P* < 0.001, \*\*\*\**P* < 0.0001 (I (Basal OCR) and K). Statistical analysis was performed using one-way ANOVA with \**P* < 0.05, \*\**P* < 0.01, \*\*\**P* < 0.001, \*\*\*\**P* < 0.0001 (F, G, I (SRC and coupling efficiency)). N represents the number of biological replicates.

Previously, it was shown that cristae shape influences the assembly of the respiratory chain complexes mediated by OPA1 (Cogliati *et al*, 2013). This led us to question whether abnormal mitochondrial ultrastructure resulting from deleting MICOS subunits could be a determinant of OXPHOS assembly specifically because certain MICOS KOs have a detrimental influence on the number of CJs and cristae while other do not. Thus, we investigated the mitochondrial ultrastructure of MICOS KOs using EM (**Fig 7E**). We found a significant loss of CJs in *MIC10*, *MIC13*, *MIC19* as well as *MIC60* KOs (**Fig 7E and F**) while a significant reduction of cristae number per unit length of mitochondria (μm) was found in *MIC10*, *MIC13* and *MIC60* KOs (**Fig 7E and G**). Thus, deletion of *MIC10, MIC13* and *MIC60* leads to stark changes in the mitochondrial ultrastructure which is also accompanied by reduced assembly of various OXPHOS complexes. Next, we checked whether reduced assembly of the OXPHOS complexes also influenced OXPHOS activity (**Fig 7H-K**). We determined mitochondrial respiration by using seahorse respirometry assays using two methods; a mitochondrial stress test determining the overall respiration and another respirometry assay determining the contribution of individual respiratory chain complexes. Despite observed assembly defects, mitochondrial respiration was largely unaffected except for coupling efficiency, which was significantly reduced in *MIC10* and *MIC60* KO cells when compared to WT HEK293 cells (**Fig 7H and I**). Checking various MICOS KO cells for the activity of complex I-, complex II- and complex IV-linked respiration by feeding corresponding substrates revealed a significant and consistent reduction in *MIC10* and *MIC60* KO cells (**Fig 7J and K**). In *MIC13* KO cells, complex II- and IV-linked respiration was significantly reduced. Overall, we reveal, using the complete set of MICOS KOs, that *MIC10* and *MIC60* KO cells showed consistent defects in OXPHOS assembly and activity. Since *MIC60* deletion leads to loss of all seven MICOS subunits uniformly while *MIC10* deletion leads to a major loss of only the MIC10 subcomplex (MIC10/MIC13/MIC26/MIC27) (Stephan *et al*., 2020), we conclude that loss of MIC10 subcomplex is sufficient to cause defects in assembly and activity of various OXPHOS complexes. In order to address the specific role of MIC10 and MIC60 in determining the OXPHOS assembly among all MICOS subunits, we also used *MIC13* KO cells which did not show a major reduction in assembly of OXPHOS complexes (except partly for complex I). For this, we employed cell lines stably expressing the pLIX403 system based on a Tet-On system. Specifically, HEK293 cells stably expressing pLIX403 with empty vector, pLIX403-MIC10, pLIX403-MIC60 or pLIX403-MIC13-Flag were used. The addition of doxycycline induces the expression of a specific gene in a temporal manner. WBs showed a successful induction of MIC10, MIC13 as well as MIC60 levels after addition of doxycycline for 24 hours validating the pLIX403 system (enclosed in solid rectangles compared to dotted rectangles, **Fig S6A**). BN-PAGE showed that the assembly defects of complex I and IV observed in *MIC10* and *MIC60* KO cells were rescued upon expression of the corresponding proteins (**Fig S6B-E**) confirming the specific role of MIC10 and MIC60 in regulating the assembly of specific OXPHOS complexes in accordance with their interaction with many proteins belonging to complex I and IV as revealed by our APEX2 proximity studies.

## Discussion

APEX is an engineered peroxidase catalysing the oxidation of biotin-phenol to generate very short-lived biotin-phenoxyl radicals (<1 ms) over a small labelling radius of <10 nm (Qin *et al*, 2021). APEX2 (single mutant version of APEX: A134P) has a greater ability to enrich proximal endogenous proteins using biotinylation and therefore is a more sensitive peroxidase compared to APEX (Lam *et al*., 2015). Accordingly, pioneering studies revealing the OM, IMS and matrix proteomes employed APEX2 instead of APEX (Hung *et al*, 2017; Hung *et al*, 2014; Rhee *et al*, 2013). The above-mentioned studies decoded the respective submitochondrial localisation by fusing APEX2 to partial sequences of specific proteins enabling their targeting to respective subcompartments. This was achieved by fusing APEX2 to C-terminal 31 amino acids of MAVS, 68 amino acid leader sequence of LACTB and mitochondrial targeting sequence of COX4I1 in order to decode the OM, IMS and matrix proteome respectively. These studies revealed a total of 137 (Hung *et al*., 2017), 127 (Hung *et al*., 2014) and 495 interaction partners (Rhee *et al*., 2013). Although increasing focus is given to proximity labelling studies using APEX2 as well as BioID2, TurboID as fusion tags (Qin *et al*., 2021), to the best of our knowledge, the interactome of any one of the two MICOS subcomplexes remained an enigma. In this endeavour, we took care to generate stable cell lines of HEK293 cells expressing MIC10-, MIC13-, MIC26- or MIC27-APEX2 fusion proteins in the background of the respective KO cell lines. Therefore, only the exogenously expressed proteins with APEX2 tags were present in these cells, and not the endogenous proteins, overcoming overexpression issues.

The MINDNet comprises 119 common and 50 different unique interactors of MIC10, MIC13, MIC26 and MIC27 and additional 20 and 44 proteins shared between any three and two MICOS subunits respectively (**Table S4**). Noteworthy, among the common interactome of the four MICOS subunits, we found all the MICOS proteins as well as MIB components SAMM50, Metaxin 1 (MTX1) and DNAJC11 (Huynen *et al*., 2016). Additionally, it is worth mentioning that few recently discovered interactors of MICOS complex were also found in our proximity labelling studies. For instance, the interaction of MYO19 with MICOS complex and SAMM50 was important for maintaining the cristae membrane architecture (Shi *et al*, 2022). MICU1 was shown to regulate cristae structure and function independent of mitochondrial calcium uniporter (MCU) (Tomar *et al*, 2023). In addition, MIC10, MIC26 and MIC60 were found in combination or alone when multiple mitochondrial ribosomal proteins and the membrane insertase OXA1 were used as baits (Singh *et al*, 2020). This study generated a mitochondrial gene expression network (MiGENet) using 40 different baits covering mitochondrial matrix proteins and is aligned with our study showing interaction of various mitochondrial ribosomal proteins with different MICOS proteins. In addition, it was recently shown that ATAD3A regulates the mitoribosome stability (Rigoni *et al*, 2025). The MINDNet also reveals ATAD3A on one hand and mitoribosomal proteins on the other hand in the proximity of MICOS subunits. Thus, the functional relationship between MICOS subunits and mitoribosomal proteins needs further investigation. Taken together, the classification of the MICOS interactome into different functional classes including OM beta barrel proteins like SAMM50, translocases of IM and OM import machinery, IM proteases including OMA1 and YME1L and CM-resident OXPHOS complexes illustrate the breadth of the interactome of the MICOS subunits. Nevertheless, it must also be taken into account that while we generated extensive proximity interaction maps of four different MICOS subunits which revealed many common as well as unique interactors, we focussed on one of the subcomplex of the MICOS proteins. Thus, it is probable that the interactome of the MIC19/MIC25/MIC60 subcomplex is at least partially distinct compared to the MIC10/MIC13/MIC26/MIC27. Here, we tried to also generate a HEK293 cell line stably expressing MIC60-APEX2 fusion protein in the background of *MIC60* KO cell lines. Despite, our best efforts, the required stable cell line could not be generated. Further, the MICOS interactome also revealed proteins possessing localisation to other organelles like ER, Golgi Apparatus, endosomes and peroxisomes indicating some leakage of the biotin-phenoxyl radical from the IMS to the mitochondrial surface via the porins in the OM. The porins allow unrestricted exchange of molecules <5 kDa between IMS and OM (Colombini, 1979). Therefore, MICOS proximity labelling revealing biotinylation of plausible interactors present at the contact sites of mitochondria with other organelles should be taken with care. Nonetheless, our proximity labelling study while revealing previously identified interactors and other MICOS subunits also showed that the MICOS interactome has a unique spatial demarcation in the IM including novel uncharacterised molecules. The localisation of the MICOS interactors in different mitochondrial subcompartments are in accordance with our expectations and also strengthen the study. Upon sorting all interactors of MICOS subunits into various groups based on the mitochondrial localisation (MitoCarta3.0 classification), we found that there is an enrichment of MICOS interactors in the OM, IMS and IM. In contrast, reduced number of MICOS interactors were present in the mitochondrial matrix. Further, the percentage of proteins sorted based on the mitochondrial functions increased for various MICOS interactors in the category of protein import and sorting machinery pathways when compared to MitoCarta3.0 classification. While 7.3 % of the total MitoCarta proteins were classified into the protein import and sorting pathway, the interactome of all four MICOS subunits with 17.9 % - 19.7 % showed a ∼2.5-fold increase. Our results are in accordance with previous literature which demonstrated that MICOS proteins interact with components of the TOMM and TIMM complex (Callegari *et al*, 2019; von der Malsburg *et al*, 2011).

An elegant BioID study involving a high-density mitochondrial proximity interactome employed MIC26 as one of the baits (Antonicka *et al*, 2020). MIC26 was found to interact with other OXPHOS proteins in line with our study. Our MINDNet data also strengthens the relationship between OXPHOS regulation and MICOS subunits. When we checked all the MICOS KOs for inefficient OXPHOS function, we found that only MIC10 and MIC60 consistently regulate the complex I and IV assembly and activity. It must be noted that although *MIC60* deletion leads to a massive loss of all MICOS proteins, depletion of the second MICOS subcomplex namely MIC10/MIC13/MIC26/MIC27 via deletion of *MIC10* is sufficient to observe OXPHOS defects. Interestingly, we found that the mitochondrial ultrastructure is only grossly affected in *MIC10*, *MIC13* and *MIC60* KO cells. Thereby, the optimal mitochondrial IM architecture via presence of CJs is necessary for determining the OXPHOS assembly and function in accordance with another study (Stoldt *et al*, 2018). Intriguingly, the CJs housing the MICOS complex were also proposed as a diffusion barrier for metabolites (Frey *et al*., 2002; Mannella *et al*., 2001). Other recent results also corroborate the relationship between MICOS and OXPHOS. Recently, it was shown, using *Saccharomyces cerevisiae*, that the IM protein Mar26 recruits Rieske Iron-sulfur protein (Rip1) to the Mic60-Mic10 module of the MICOS complex to coordinate the assembly of OXPHOS complex III (Zerbes *et al*, 2024). This study involving human cell lines also revealed TTC19 as a common MICOS interactor. Previously, TTC19 was shown to be a regulator of complex III biogenesis (Bottani *et al*, 2017). TTC19 helps to remove the N-terminal fragments of the Rieske protein UQCRSF1, which inhibits complex III maturation. MIC19 was also found to associate with COX subunit IV (Sastri *et al*, 2017). Intriguingly, while our proximity labelling study revealed the majority of enriched complex IV subunits to be common, we also found certain proteins in the proximity neighbourhood of only MIC10 and MIC13 but not MIC26 and MIC27. MT-CO3 and COX4I1 were found to be in the vicinity of both MIC10 and MIC13. In accordance, we found stark defects in complex IV assembly and activity in *MIC10* and *MIC13* KO cells. Thus, spatial demarcation of complex IV protein, mt-CO3 in the neighbourhood of MIC10 and MIC13 and not MIC26 and MIC27 reinforce the role of MIC10 and MIC13 in regulating complex IV function. Overall, our study links the MICOS complex to complex IV assembly and activity in mammalian cells by revealing specific protein-protein proximities.

We could envisage that the MICOS subunits strategically positioned at the CJs interact with important proteins belonging to SLC family in order to integrate mitochondrial IM architecture and metabolic homeostasis. In essence, while our study underlines the breadth of the MICOS interactome at steady-state, it remains to be determined how the interactome undergoes dynamic changes, under conditions of altered nutrient availability as well as in the background of different deletion mutants, defining a broad range of physiological processes covering diverse mitochondrial functions. In other words, the MINDNet provides a foundation for future studies which aim to elucidate the dynamic IM interactome covering different subcompartments within mitochondria.

## Methods

### Cell culture

Flp-In T-REx HEK293 cells were grown in 1 g/L glucose DMEM (PAN-Biotech) along with the following supplements: 10 % FBS (Capricorn Scientific), 2 mM stable glutamine (PAN-Biotech) and penstrep (PAN-Biotech, penicillin 100 U/mL and 100 μg/mL streptomycin). DMEM high glucose medium (PAN-Biotech, P04-03500) supplemented with 10 % FBS, 2 mM stable glutamine, 1 mM sodium pyruvate (Gibco, 11360070) was used for culturing of Plat-E and HEK293FT cells. Cells were cultured at 37 °C supplied with 5 % CO_2_.

### CRISPR-Cas9 knockout generation

Individual transfection of the respective CRISPR-Cas9 double nickase plasmids (Santa Cruz Biotechnology, MIC26: sc-413137-NIC, MIC60: sc-403617-NIC, MIC25: sc-413621-NIC, MIC19: sc-408682-NIC, MIC10: sc-417564-NIC, MIC27: sc-414464-NIC) was performed with GeneJuice (Sigma-Aldrich, 70967-3) in Flp-In T-REx HEK293 WT cell line to generate the corresponding MICOS KO cell lines. The KO cell lines were already employed in one of our recent previous study (Naha *et al*., 2024). Briefly, cells were transfected at 60-70 % confluency with 1 µg of double nickase plasmid for 48 h followed by 2.5 µg/ml puromycin selection for 24 h. Subsequently, based on green fluorescent protein (GFP) expression, single cell sorting was performed using flow cytometry in 96-well plates. Cell population was expanded and screened for successful KO generation using WBs. Cell lines showing no immune reactivity to respective antibodies were considered as KO cell lines.

### Molecular cloning

Human ORFs for *MIC10*, *MIC13*, *MIC26* and *MIC27* were cloned along with APEX2 (Addgene, 49386) into pMSCVpuro vector using Gibson assembly cloning kit (NEB, E2611L) following the manufacturer’s protocol. For generating the IM-APEX2, the initial 116 amino acids of the mitoT plasmid (Viana *et al*., 2021) was used to amplify the mitochondrial signal peptide followed by the IM transmembrane domain. MIC10 and MIC60 ORFs were cloned into pLIX403 (Addgene, 41395) with Gibson assembly according to manufacturer’s instructions. Primer sequences for Gibson assembly cloning are provided in the key resources table.

### Generation of stable cell lines

For retroviral transduction, Plat-E cells (Anand *et al*, 2020) were co-transfected with 1 µg of pMSCV-MIC10/MIC13/MIC26/MIC27-APEX2 or pMSCV-IM-APEX2 and 1 µg of pVSV-G along with 3.5 µL of GeneJuice reagent per 6-well plate. After 72 h incubation, viral supernatant was collected and added to the target HEK293 cells.

For lentiviral transduction, HEK293FT cells were transfected with 1 µg of pLIX403 EV or pLIX403-MIC13-Flag, pLIX403-MIC10 or pLIX403-MIC60 along with 1 µg of psPAX2 (Addgene, 12260) and pMD2.G (Addgene, 12259) using GeneJuice transfection reagent. 72 h post transfection, viral supernatant was added on target cell lines.

In both cases, puromycin selection (2.5 µg/ml) was initiated 48 h post-transduction and carried out for 2 weeks in total. The expression of exogenous proteins was confirmed with WBs.

### SDS-PAGE and Western Blotting

Cells were harvested by scraping after three washes with DPBS (PAN-Biotech). After pelleting (1000 g, 5 min, 4 °C), cells were resuspended in an appropriate volume of RIPA buffer (150 mM NaCl, 0.1 % SDS, 0.05 % Sodium deoxycholate, 1 % Triton-X-100, 1 mM EDTA, 1mM Tris, pH 7.4, 1x protease inhibitor (Sigma-Aldrich)) and incubated for 30 min on ice. The DC^TM^ protein assay kit (BIO-RAD, 5000116) was used to determine the protein concentration. SDS samples were prepared in Laemmli buffer followed by heating at 95 °C for 5 min. A range of SDS electrophoresis gels (8 %, 10 %, 12 % or 15 %) were used for separating samples based on the molecular weights of individual proteins. WB was performed using nitrocellulose membranes and transfer efficiency was verified using Ponceau S (Sigma Aldrich). Following destaining, nitrocellulose membranes were blocked using 5 % milk in TBS-T for 1 h at room temperature (RT), probed at 4 °C overnight with 1:1000 dilutions of primary antibodies in 5 % milk in TBS-T. The following HRP conjugates of secondary antibodies were used: Goat IgG anti-Rabbit IgG (Dianova, 1:10000) and Goat IgG anti-Mouse IgG (Abcam, 1:10000) conjugated to HRP. Chemiluminescence was recorded using Signal Fire ECL reagent (Cell Signaling Technology) and VILBER LOURMAT Fusion SL equipment (Peqlab).

### APEX2 proximity biotinylation

Stable Flp-In T-REx HEK293 cell lines expressing MICOS-APEX2, in the background of respective MICOS KO cell lines, were used for proximity biotinylation experiments. WT Flp-In T-REx HEK293 cell lines expressing IM-APEX2 stably and matrix-APEX2 transiently were used as control cell lines. Biotin-phenol labeling was performed by pre-incubating the cells with 0.5 mM biotin-phenol for 30 min. To start the reaction, 0.5 mM H_2_O_2_ in DPBS was added to the cells and incubated for 1 min at RT. The reactions were quenched by 3 washes with 10 mM sodium azide, 10 mM sodium ascorbate, and 5 mM Trolox in DPBS. The cells were recovered and solubilized with RIPA Buffer containing 10 mM sodium ascorbate, 10 mM sodium azide, 5 mM Trolox, and protease inhibitor. Prior to mass spectrometry (MS) analysis, reaction efficiency was assessed by SDS-PAGE and WBs using streptavidin-HRP (1:1000, Merck).

Streptavidin-magnetic beads (Thermo Scientific) were used to purify APEX2-biotinylated proteins. 500 μg of total cell lysates containing biotinylated proteins were used and processed for MS analysis. The beads were equilibrated with RIPA buffer containing 10 mM sodium ascorbate, 10 mM sodium azide, 5 mM Trolox, and protease inhibitor. Samples were loaded overnight at 4 °C. Next morning beads were washed three times with RIPA buffer followed by three more washes with ABC buffer. Samples were denatured with 50 µl of urea buffer (6 M urea, 2 M thiourea) and followed by disulfide-bridge reduction using dithiothreitol at a final concentration of 5 mM for 1 h at RT. To alkylate oxidized cysteines, 2-Chloroacetamide was added to the samples until a concentration of 40 mM was reached and incubated for 30 min in the dark. Samples were finally diluted 1:3 with ABC buffer to reach 2 M urea concentration. Protein digestion was performed overnight with trypsin (1:100). Following acidification with 1 % formic acid, samples were desalted using a modified version of the Stop and Go extraction tip (StageTip) protocol. For that, the acidified samples were centrifuged at full speed in a table top centrifuge for 5 min and the supernatants were loaded onto the equilibrated StageTips by centrifugation at 2600 rpm for 5 min. StageTips equilibration was performed by subsequent washes with methanol, 0.1 % formic acid in 80 % acetonitrile and 2 washes with 0.1 % formic acid in water. All equilibration steps were followed by centrifugation at 2600 rpm for 1-2 min in order to dry the StageTip membrane. After loading the sample, the StageTip was washed with 0.1 % formic acid in water and centrifuged at 2600 rpm for 3 min, followed by two washes with 0.1 % formic acid in 80 % acetonitrile with subsequent centrifugation at 2600 rpm for 3 min. Finally, the StageTips were dried using a syringe and stored at 4 °C until further processing.

### Proteomics data acquisition

Samples were analyzed on an Orbitrap Exploris 480 mass spectrometer (Thermo Fisher Scientific). Samples were loaded onto a precolumn (Acclaim 5µm PepMap 300 µ Cartridge) with a flow of 60 µl/min before reverse-flushing onto an in-house packed analytical column (30 cm length, 75 µm inner diameter, filled with 2.7 µm Poroshell EC120 C18, Agilent). Peptides were chromatographically separated with an initial flow rate of 400 nl/min and the following gradient: initial 2 % solvent B (0.1 % formic acid in 80 % acetonitrile), up to 6 % in 3 min. Then, flow was reduced to 300 nl/min and solvent B increased to 20 % in 26 min, up to 35 % B within 15 min and up to 98 % solvent B within 1.0 min while again increasing the flow to 400 nl/min, followed by column wash with 95 % solvent B and re-equilibration to initial condition. The mass spectrometer was operated in data-dependent acquisition with a cycle time of 1 s with MS1 scans acquired from 350 m/z to 1400 m/z at 60k resolution and an AGC target of 300 %. MS2 scans were acquired at 15 k resolution with a maximum injection time of 118 ms and a normalized AGC target of 50 % in a 2 Th window and a fixed first mass of 110 m/z. All MS1 scans were stored as profile, all MS2 scans as centroid. Raw files were processed with Maxquant (version 2.4, (Tyanova *et al*, 2016)) using the human UniProt database (UP000005640) and default parameters with the match-between-runs option enabled between replicates. The mass spectrometry data have been deposited to the ProteomeXchange Consortium via the PRIDE partner repository (Perez-Riverol *et al*, 2022) with the dataset identifier PXD063925.

### Bioinformatic analysis and software packages

The Bioinformatics data analysis was performed using the R Statistical Software (R version 4.4.2., 2024-10-31) (R Core Team, 2024) including few Bioconductor packages (Huber *et al*, 2015). Log_2_ transformation and median normalization were applied on the complete dataset. PCA analysis was done and a visual representation created using the package factoextra (Kassambara & Mundt, 2020). Statistical testing to identify differentially abundant proteins was performed with the package limma (Ritchie *et al*, 2015). Volcano Plots were generated using the package Enhanced Volcano (Blighe *et al*, 2024). Database information of MitoCarta3.0 (Rath *et al*., 2021) was used to filter mitochondrial proteins/genes and to retrieve reliable information about pathway and mitochondrial localization. In addition, Cytoscape (Shannon *et al*, 2003) and Venny (Oliveros, 2007-2015) were used to make figures in the manuscript where required. The MICOS interactors using the log_2_FC enriched conditions were considered when the proteins were present in at least three of the five replicates in both control as well as MICOS-APEX2 conditions. For the FOM (**F**ound **O**nly in respective **M**ICOS-APEX2) enrichment conditions, the proteins were present in at least one of the respective MICOS-APEX2 replicate and absent in the matrix-APEX2 or IM-APEX2 control.

### Blue Native-PAGE

15 cm dishes were used to seed 3 x 10^6^ Flp-In T-REx HEK293 and grow until approx. 80 % confluency. Cells were washed with cold PBS buffer, scraped and pelleted at 1000 g, 4 °C for 5 min. 1 ml of lysis buffer (210 mM mannitol, 70 mM sucrose, 1 mM EDTA, 20 mM HEPES, 0.1 % BSA, 1 x protease inhibitor) was used to resuspend the cell pellets. After 10 min incubation on ice, cells were lysed by mechanical disruption with repetitive strokes employing a 20G canula and differential centrifugation steps at 1000 x g, 4 °C for 10 min to remove heavy membranes and 10,000 x g, 4 °C for 15 min to pellet mitochondria. Protein concentration was measured with a DC Protein Assay Kit from the mitochondrial suspension in BSA-free lysis buffer.

For performing the blue native PAGE, 2.5 g/g digitonin to protein ratio was used to solubilize 100 µg of mitochondria. After 1 h of incubation on ice, the samples were centrifuged for 20 min at 20,000 x g and 4 °C. The supernatants were supplemented with loading buffer (50 % glycerol, 8 g/g Coomassie to detergent ratio) and promptly loaded onto 3-13 % gradient gel. Complexes were separated using 150 V and 15 mA for 16 h. Then, Western blot transfer was performed using PVDF membranes, which were destained with methanol and blocked overnight at 4 °C, with 5 % milk in TBS-T. For identification of relevant protein complexes, the membranes were probed using the following antibodies: NDUFB4 (Abcam, 1:1000), UQCRC2 (Abcam, 1:1000), COXIV (Abcam, 1:1000), ATP5A (Abcam, 1:1000), Goat IgG anti-Mouse IgG (Abcam, 1:10000) and Goat IgG anti-Rabbit IgG (Dianova, 1:10000) conjugated to HRP. Pierce™ SuperSignal™ West Pico PLUS chemiluminescent substrate reagent (Thermo Scientific) and VILBER LOURMAT Fusion SL equipment (Peqlab) were used to obtain the chemiluminescent signals.

### Electron Microscopy

Flp-In T-REx HEK293 cells (4 x 10^6^) were grown for 48 h in 10 cm petri dishes at 37 °C with 5 % CO_2_. Cells were fixed using 3 % glutaraldehyde in 0.1 M sodium cacodylate buffer (pH 7.2). For staining the APEX2 activity, cells were incubated for 10 min in 20 mM glycine in 0.1 M sodium cacodylate buffer. After 3 washes with sodium cacodylate buffer, cells were pre-incubated with 1,4 mM 3,3’-Diaminobenzidine (DAB) tetrahydrochloride hydrate in cacodylate buffer for 15 min followed by a 20 min incubation in fresh DAB solution with 10 mM H_2_O_2_. The reaction was terminated by changing the reaction solution against cacodylate buffer and cells were processed following the standard EM sample preparation protocol. Cell pellets were washed with 0.1 M sodium cacodylate and submerged in 2 % agarose. Contrasting was performed with 1 % osmium tetroxide for 50 min, washed twice with sodium cacodylate and incubated in 1 % uranyl acetate/1 % phosphotungstic acid for 1 h. Sample dehydration was performed with graded acetone series and polymerization was performed via embedding in spur epoxy resin at 65 °C for 24 h. Ultrathin sections of samples were prepared with microtome and images were captured with transmission electron microscope (Hitachi, H7100) at 100kV equipped with Morada camera or with JEOL JEM-2100plus (80 kV or 200 kV) with Matataki flash camera and analyzed with ImageJ software. The images were randomized and the data was analyzed in a double-blind manner by independent scientists. Data analysis was carried out by GraphPad prism.

### Confocal and STED super-resolution microscopy

For sample preparation, Flp-In T-REx HEK293 cells were seeded onto 35 mm GelTrex-coated dishes and cultured until 70-80 % confluency. Cells were fixed using 4 % paraformaldehyde (Sigma-Aldrich, P6148) for 20 min at RT and washed 3x with DPBS followed by subsequent permeabilization with 0.15 % Triton X-100 (Sigma-Aldrich, T8787) at RT for 15 min. Permeabilized cells were washed once with DPBS and treated with blocking solution (1 % goat serum) for 15 min at RT. Further, the blocking solution was removed and primary antibodies (1:100) were added onto the coverslips and incubated at 4 °C overnight. The following antibodies were used for confocal microscopy: MIC60 (Abcam, ab110329), Streptavidin DyLight 488 (Thermo Fisher, 21832), Goat anti-mouse Alexa Fluor 646 was used for binding the MIC60 primary antibody. For preparing the samples for STED SR imaging, we used the following secondary antibodies: Goat anti-mouse Aberrior STAR 635P and Goat anti-rabbit Aberrior STAR 580 (both 1:100). For marking the biotin, we used an anti-biotin primary antibody (1:100).

Confocal microscopy was performed using a spinning disc confocal microscope (Perkin Elmer) equipped with a 60x oil-immersion objective (N.A = 1.49) and a Hamamatsu C9100 camera (1000 X 1000 pixel). The images were obtained using 488 nm and 640 nm excitation laser lines and 527 nm (W55) and 705 nm (W90) emission lines respectively. Dual-colour STED SR imaging was performed using Leica SP8 laser scanning confocal microscope coupled with a STED module using a 93x glycerol objective (N.A = 1.3). White light laser excitation wavelengths of 562 nm and 633 nm were employed to detect the emission signals between 571-628 nm and 643-699 nm respectively using hybrid detectors (HyDs) with a pinhole of 1 AU. For both the channels, a pulsed STED depletion laser beam of 775 nm wavelength was used. In order to optimise the specificity of the signal, Gating STED was used from 1.2 ns onwards. The channels were acquired sequentially with a field of view of 9.62 x 9.62 µm achieved at 13x magnification. For image depiction, smoothing was carried out with Fiji software and the contrast was adjusted for optimal visibility. Before imaging, the excitation and depletion laser alignments were checked using reflection mode with the help of colloidal 80 nm gold particles (BBI Solutions). Chromatic aberration between both channels was ruled out by using an anti-Nup153 antibody emitting in two channels.

### Mitochondrial Respirometry

Respirometry experiments were conducted using Seahorse XFe96 Analyzer (Agilent). Flp-In TREx HEK293 cells were seeded onto Poly-D-Lysine-coated (50 µg/mL) Seahorse plates (4×10^4^ cells/well) and grown in normal cell culture medium overnight at 37 °C and 5 % CO_2_. For performing the mitochondrial stress test, cells were washed twice with Seahorse assay media supplemented with 10 mM glucose, 2 mM stable glutamine and 1 mM sodium pyruvate and incubated in 180 µl fresh Seahorse assay medium with the above-mentioned additives for 45 min at 37 °C in CO_2_-free incubator prior to the assay. Mitochondrial oxygen consumption was measured after sequential addition of 1 µM oligomycin (Sigma), 0.5 µM FCCP (Sigma) and 0.5 µM rotenone/antimycin a (Sigma) using the manufacturer’s instructions.

For the assessment of individual OXPHOS complex activities, cells were washed twice with 1x MAS buffer (2 mM HEPES, 220 mM mannitol, 70 mM sucrose, 10 mM KH2PO4, 5 mM MgCl2, 1 mM EGTA, 0.2 % BSA) and then supplied with fresh MAS buffer supplemented with 1 nM PMP (Agilent), 1 mM malate, 10 mM pyruvate and 4 mM ADP following which the plate was immediately placed in Seahorse XFe96 Analyzer and the assay initiated. Oxygen consumption was assessed after sequential injections of 2 µM rotenone, 10 mM succinate, 2 µM antimycin and 10 mM ascorbate with 200 µM TMPD. In order to reduce the assay time, due to the exposure of cells to a permeabilizing agent, mixing and waiting times were restricted to 30 s and measurement time limited to 2 min.

For normalization assay for cell number, medium was carefully removed from the wells using a multi-channel pipet and 100 μl of 10 μg/ml Hoechst solution (Thermo Fisher 62249) was added to the individual wells. After 10 min incubation in the dark, fluorescence was measured at 486 nm using a TECAN plate reader.

### Statistics and data representation

Data are represented by using mean ± standard error mean (SEM). Statistical significance was determined by one-way ANOVA or one sample *t*-test and mentioned in the respective figure legends with \**P* < 0.05, \*\**P* < 0.01, \*\*\**P* < 0.001, \*\*\*\**P* < 0.0001. For MS data, statistical testing was performed with the R package limma using Benjamini-Hochberg adjusted *P*-values for the significance tests.

## Key resources table

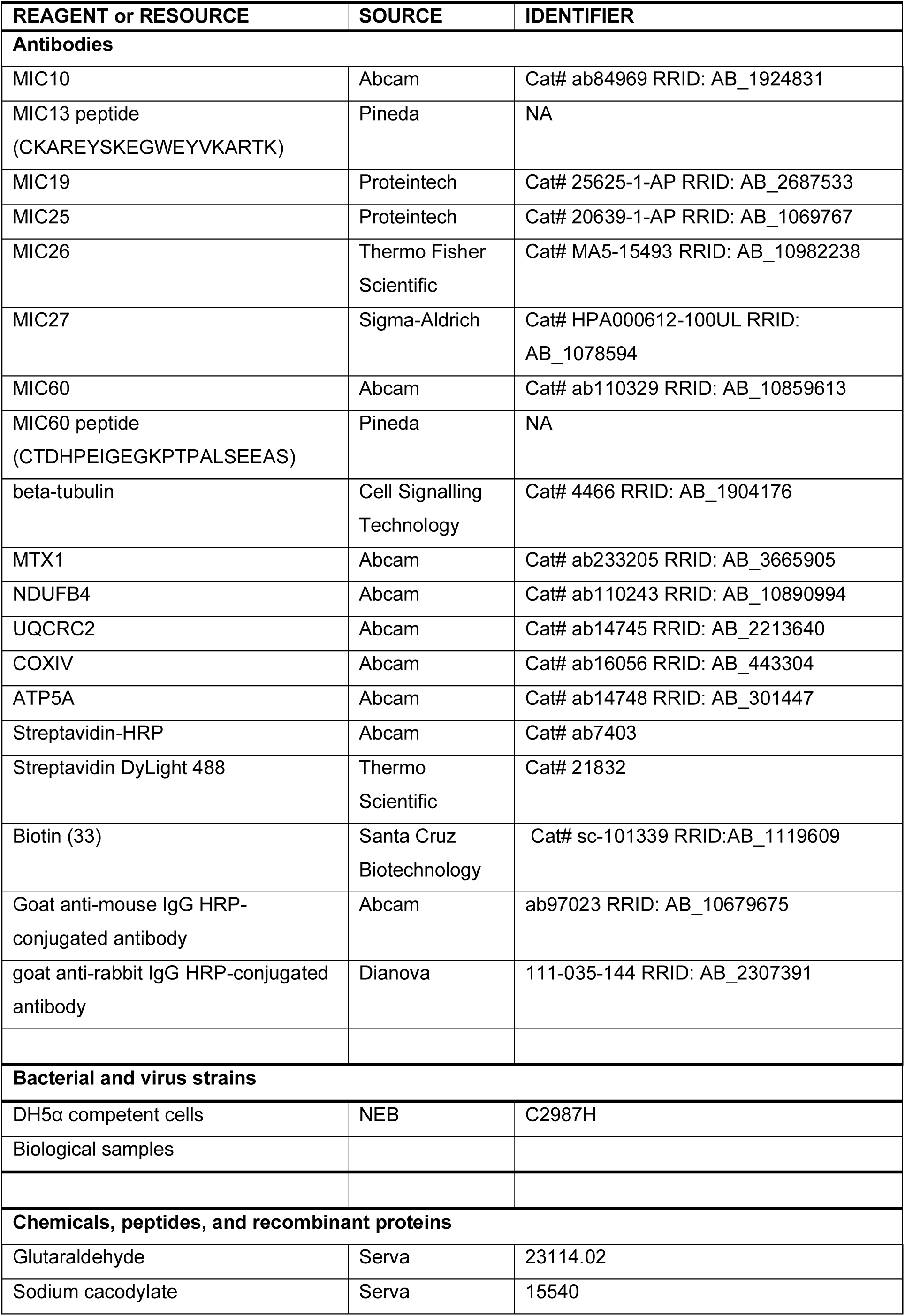
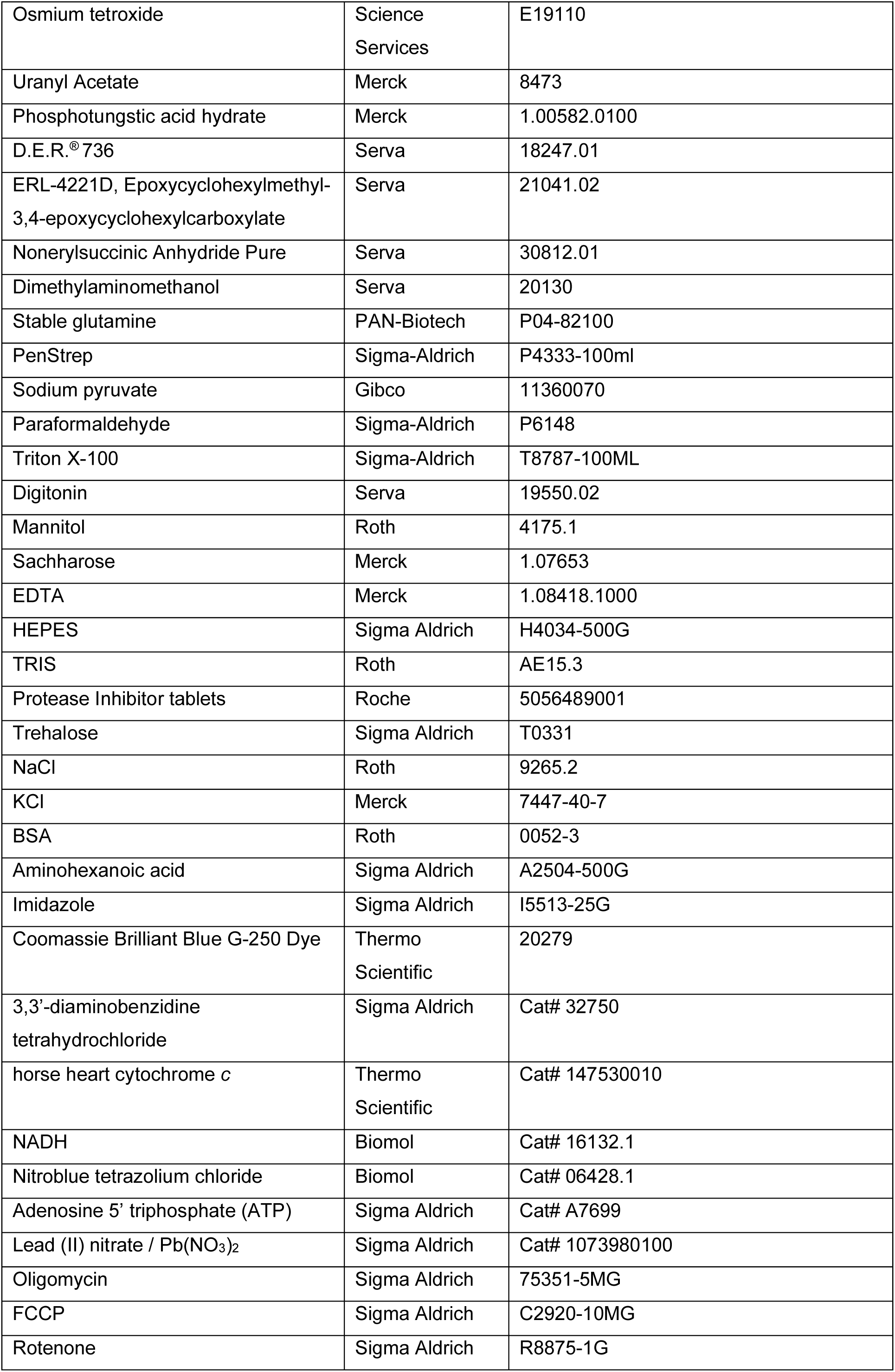
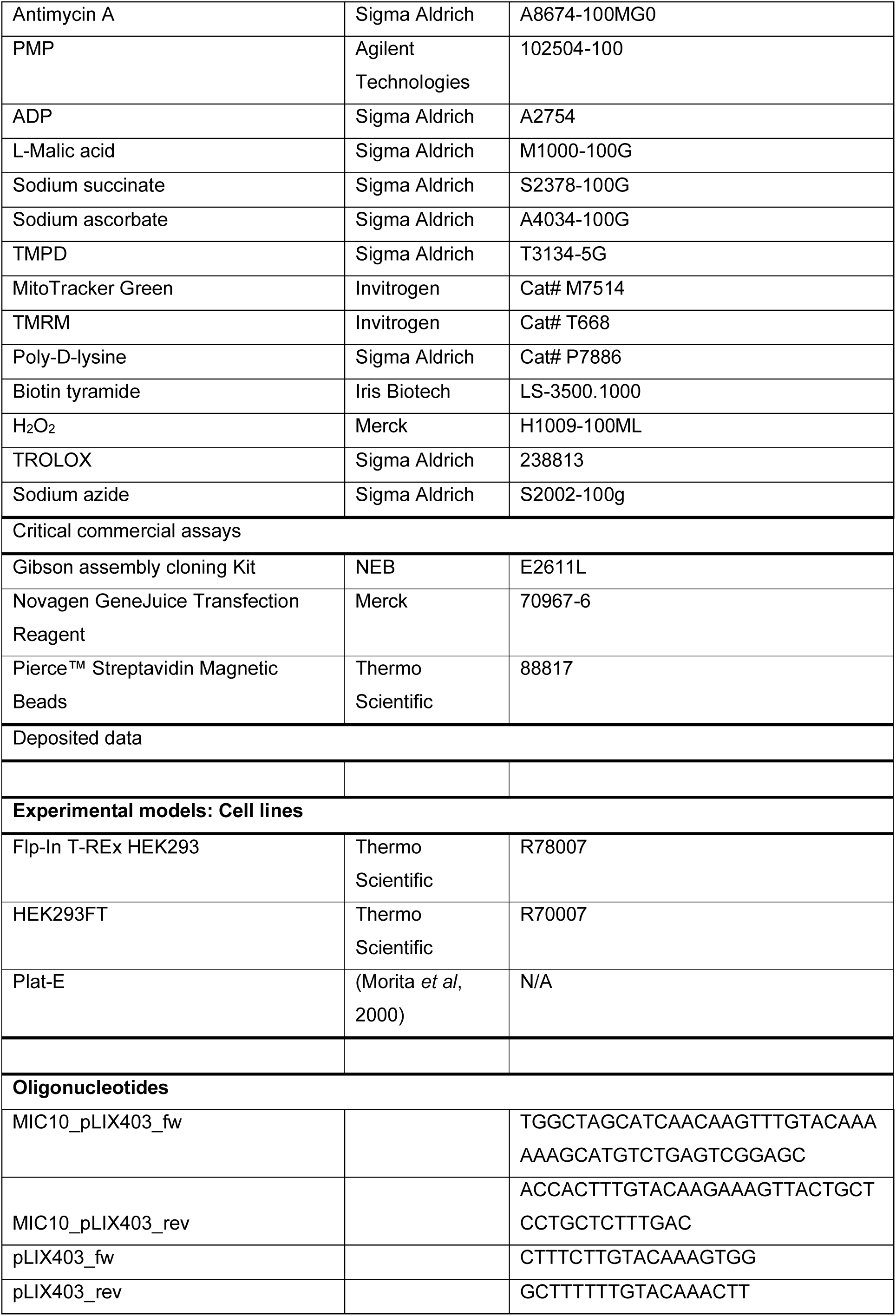
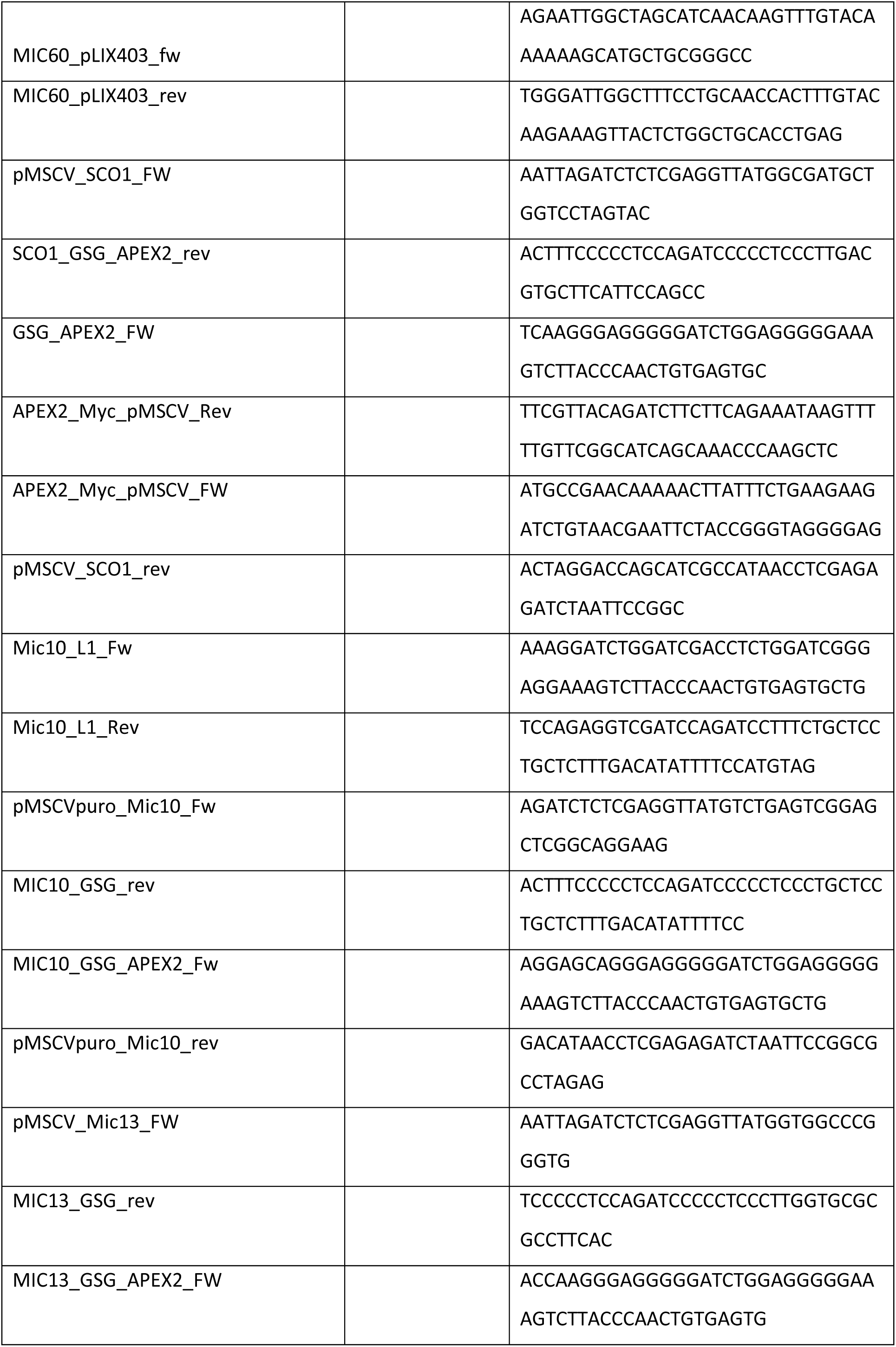
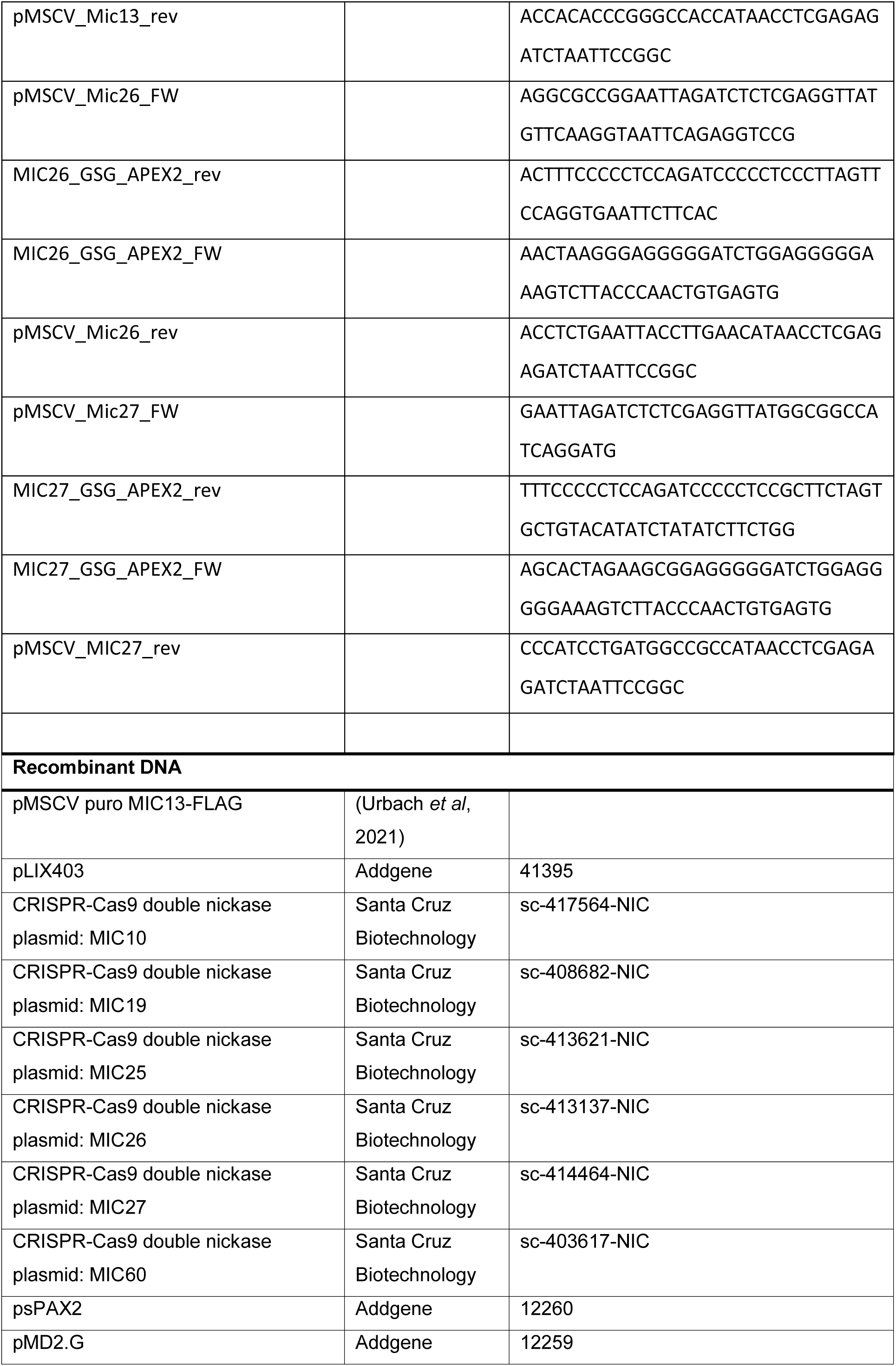
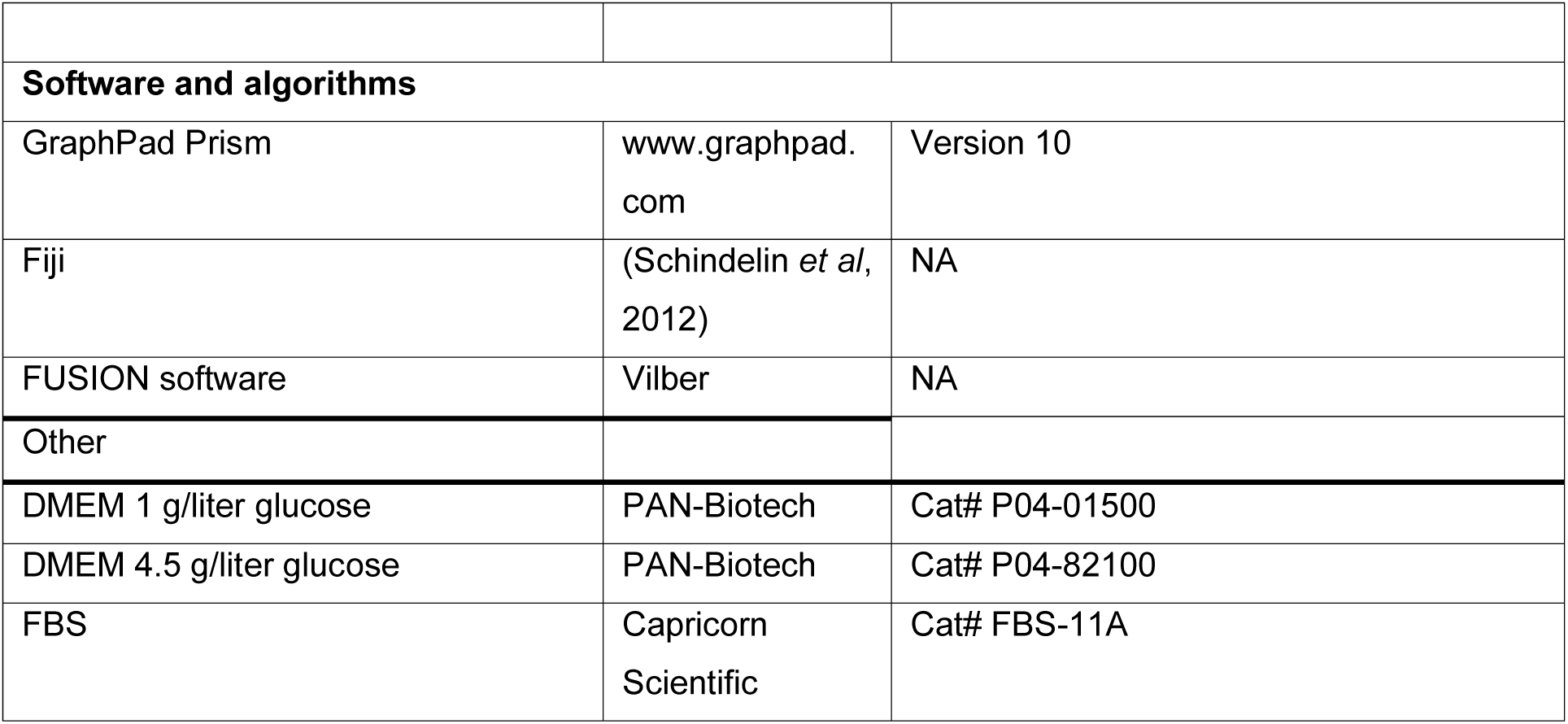

## Author Contributions

Y Schaumkessel: Investigation, methodology, visualization, validation, data curation, formal analysis and writing–original draft, review and editing.

N Brocke-Ahmadinejad: Investigation, methodology and formal analysis.

R Strohm: Investigation, methodology and formal analysis.

S Müller: Resources, investigation and formal analysis.

AS Reichert: Funding acquisition, project administration, resources and writing–review and editing.

AK Kondadi: Conceptualization, Funding acquisition, project administration, supervision, visualization, resources and writing–original draft, review and editing.

## Acknowledgements

The project was funded by the German Research Foundation (Deutsche Forschungsgemeinschaft (DFG) through the SFB 1208 project B12 (ID 267205415) to A.S.R and via the DFG Grant KO 6519/1-1 to A.K.K. We are grateful to Tanja Portugall, Andrea Borchardt and Gisela Pansegrau for providing technical assistance with molecular cloning, electron microscopy and cell culture experiments, respectively. We are thankful to Ruchika Anand for constructive discussions and David Pla-Martin and Oleh Khalimonchuk for sharing the Matrix-APEX2 and MitoT plasmids, respectively. We thank Melissa Damiecki for help with skeletonization of the EM data. Electron microscopy experiments were performed at the core facility for electron microscopy (CFEM) of the medical faculty, Heinrich Heine University Düsseldorf. We thank the CECAD Proteomics facility for the performing the experiments and analysis of proteome data and Center for Advanced Imaging (CAi), HHU where the STED super-resolution experiments were performed. This work was supported by the large instrument grant INST 216/1163-1 FUGG and INST 208/755-1 FUGG by the DFG major research equipment application (Großgeräteantrag). Alphafold 3 was used to generate 3D molecular structures predictions while PyMOL (Version 2.5.7) was used for visualization and pseudocolouring of the molecular structures. Many thanks to Athanasios Papadopoulos from Center for Structural Studies (CSS) HHU for the help with the export of visualized structures.

## Declaration of interests

The authors declare no competing interests

## Supplementary Information

**Figure S1. Expression and catalytic activity of the APEX2 fusion proteins**

(A) Western blot (WB) analysis showing the tested titration ranges of biotin-phenol (BP) in WT HEK293 cells expressing IM-APEX2. A concentration of 0.5 mM biotin-phenol was optimal for biotinylation in the mitochondrial IM.

(B) WB analysis, showing the tested H_2_O_2_ titration ranges with 0.5 mM BP in WT HEK293 cells expressing IM-APEX2, demonstrate the optimal concentration of 0.5 mM H_2_O_2_ for biotinylation experiments.

(C) WB analysis showing the distinct biotinylation pattern of HEK293 cells expressing various MICOS-APEX2 proteins and matrix-APEX2 control upon employing optimal BP (0.5 mM) and H_2_O_2_ (0.5 mM) concentration suitable for proximity labeling.

**Figure S2. Principal component analysis plots and overview of proteins present in the number of biological replicates in mass spectrometry analysis**

(A and B) Principal component analysis (PCA) plot, depicting the clustering of individual samples subjected to mass spectrometry, display a very close correlation between the MICOS-APEX2 fusion proteins. The matrix-APEX2 (A) manifest as distinct clusters from all the four MICOS-APEX2 fusion proteins, whereas the IM-APEX2 (B) group shows a similar distribution as the MICOS-APEX2 groups.

(C and D) Representation of the respective number of proteins detected in the corresponding number of technical replicates in MICOS-APEX2, IM-APEX2 and matrix-APEX2 controls covering the whole cell (C) and mitochondria (D).

**Figure S3. Flow chart and analysis pipeline of the interactome of all the four MICOS-APEX2 fusion proteins**

(A – C) Flow-chart of the analysis pipeline consisting of multiple filtering steps. The initial data set, containing 4,245 proteins, was reduced to 4,119 proteins after removal of reverse and potential contaminants. A maximum of 2613 to 2892 proteins were detected in all the four MICOS interactomes. Overall, 1146 to 1606 proteins were enriched including the log_2_FC enriched (increase ≥ 0) and FOM category (normalized to both the controls). The individual distribution of enriched proteins into different categories is shown (Fig 3A Step1, Fig 3B and C histograms). Post MitoCarta3.0 filtering, the number of proteins for various MICOS interactomes is shown (Step 2). Further, the enriched proteins after using log_2_FC cut-off of ≥ 1 for matrix-APEX2 and log_2_FC ≥ 0.5 for IM-APEX2 are shown (Step 3). Note the number of enriched proteins belonging to the FOM category (where no proteins were found in respective controls) does not change in step 3.

(D) Pie charts depicting the fractions of mitochondrial and non-mitochondrial proteins present in the interactome of respective MICOS proteins after the application of log_2_FC cut-offs of ≥1 or ≥ 0.5 for matrix-APEX2 and IM-APEX2 controls, respectively, and including the FOM candidates.

**Figure S4. Volcano plots showing the MICOS interactors (Log_2_FC enriched category)**

(A – H) Volcano Plots showing the enriched MIC10-APEX2 (A), MIC13-APEX2 (B), MIC26-APEX2 (C), MIC27-APEX2 (D) proximity proteome when normalized to the matrix-APEX2 control as well as the IM-APEX2 control (E, F, G and H) after MitoCarta3.0 filtering. The top 10 hits with the highest log_2_FC enrichment are shown. OPA1 was the most enriched in all the four interactomes when compared to matrix-APEX2 control.

**Figure S5. Expression of various proteins in individual MICOS HEK293 KO cells**

(A and B) Representative WB analyses of various MICOS subunits in the individual MICOS KO cell lines (A), along with the quantification from various biological replicates, reveals that deletion of either *MIC10*, *MIC13*, *MIC19* or *MIC60* results in total loss or strong reduction of all other MICOS subunits. Deletion of *MIC25*, *MIC26* or *MIC27* has no effect on other MICOS subunits.

Data are represented as mean ± SEM. Statistical analysis was performed using one sample *t*-test with \**P* < 0.05, \*\**P* < 0.01, ***P < 0.001, \*\*\*\**P* < 0.0001.

**Figure S6. Demonstration of specificity of MIC10 and MIC60 in regulating the assembly of OXPHOS complexes I and IV**

(A) WB analysis, showing the stable expression of either pLIX403-MIC10, pLIX403-MIC60 or pLIX403-MIC13-Flag along with pLIX403 empty vector control, after addition of doxycycline using the Tet-On system. Absence of MICOS proteins in corresponding MICOS KO cell lines is shown by dotted rectangles while the recovery is show using solid rectangles after doxycycline addition.

(B-E) BN-PAGE analyses of various OXPHOS complexes before and after doxycycline addition in the mentioned MICOS KO cell lines. A specific increase of OXPHOS complex I (B) and IV assembly (D) were observed after MIC10 and MIC60 induction in the Tet-On system. The rescue of OXPHOS complex I and IV assembly is shown using solid rectangles in *MIC10* and *MIC60* KO cells upon comparison before doxycycline addition (dotted rectangles).

## Supplementary Tables

**Supplementary Table S1**

Raw data of label-free mass spectrometry analysis of the biotinylated proteins after APEX2 labeling reaction and streptavidin pulldown. Columns comprise protein identifiers, IBAQ values, razor and unique peptides as well as MS/MS counts for the global analysis. Normalized maxLFQ (label-free quantitation) and number of peptides found in the individual technical replicates, along with the information on protein identification such as sequence coverage, identification type and peptide sequences detected in the different APEX2 groups, matrix-APEX2 and IM-APEX2 controls.

**Supplementary Table S2**

Data set comprising log_2_FC and Benjamini-Hochberg adjusted *P*-values along with the number of technical replicates in which the protein was detected for the comparison of the respective MICOS-APEX2 groups to the matrix-APEX2 control. For Log_2_FC candidates, the requirement for detection frequency of at least three technical replicates had to be met (additional columns with binary information on the fulfillment of this requirement included). All candidates not detected in the respective control, but detected in at least one technical replicate in MICOS-APEX2 were considered as FOM candidates. MitoCarta3.0 sorted hits and log_2_ median normalized LFQs of the individual replicates of the respective groups are enclosed in sheet 2 and 3, respectively.

**Supplementary Table S3**

Data set comprising log_2_FC and Benjamini-Hochberg adjusted *P*-values along with the number of technical replicates in which the protein was detected for the comparison of the respective MICOS-APEX2 groups to the IM-APEX2 control. For log_2_FC candidates, the requirement for detection frequency of at least three technical replicates had to be met (additional columns with binary information on the fulfillment of this requirement included). All candidates not detected in the respective control, but detected in at least one technical replicate in MICOS-APEX2 were considered as FOM candidates. MitoCarta3.0 sorted hits and log_2_ median normalized LFQs of the individual replicates of the respective groups are shown in sheet 2 and 3, respectively.

**Supplementary Table S4**

Summarizing overview for the enriched proteins in individual MICOS groups including log_2_FC-thresheld (*P*-value ≤ 0.05 and log_2_FC ≥ 1 or log_2_FC ≥ 0.5, for Matrix-APEX2 or IM-APEX2, respectively) and FOM candidates along with their localization and function as assigned in MitoCarta3.0. An overview of the MINDNet candidates including relevant log_2_FC values and FOM candidates (indicated with “+” in the respective cell lines) is enclosed in sheet 2.

## Notes

### Competing Interest Statement

The authors have declared no competing interest.

