## Supplementary Figures for "MINDNet: Proximity interactome of the MICOS complex revealing a multifaceted network orchestrating mitochondrial biogenesis"

#### Supplementary Figure 1

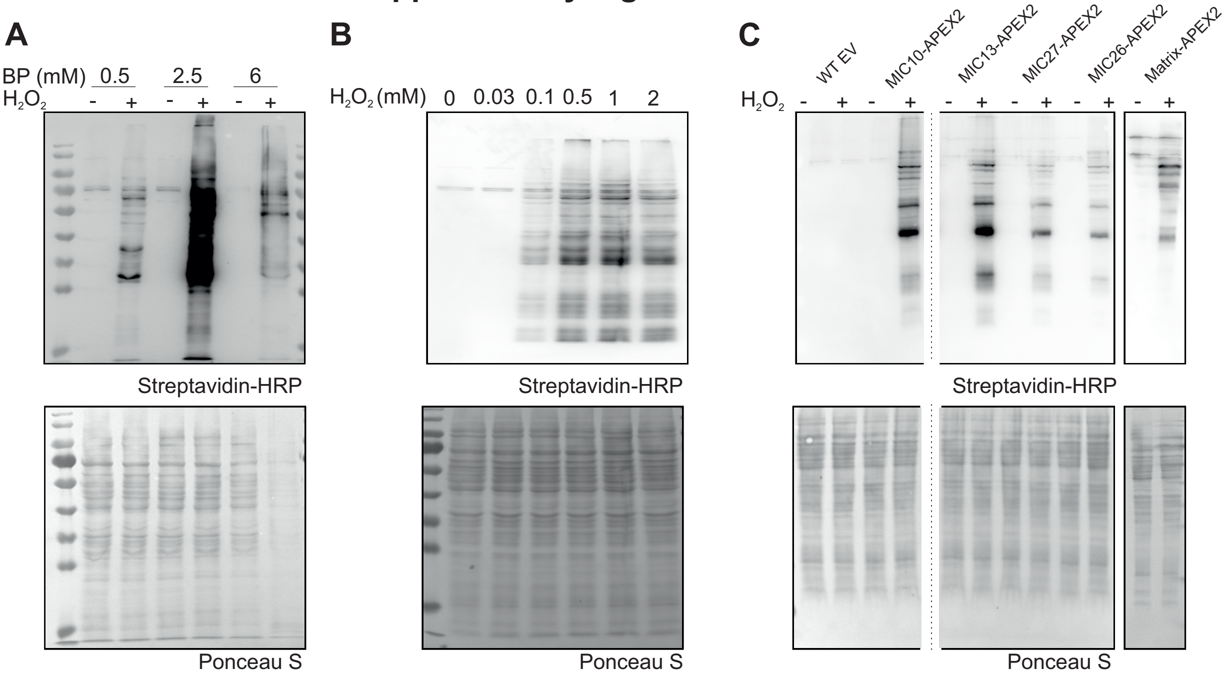


**Figure S1:** Expression and catalytic activity of the APEX2 fusion proteins

**Figure S1:** Expression and catalytic activity of the APEX2 fusion proteins

(A) Western blot (WB) analysis showing the tested titration ranges of biotin-phenol (BP) in WT HEK293 cells expressing IM-APEX2. A concentration of 0.5 mM biotin-phenol was optimal for biotinylation in the mitochondrial IM.

(B) WB analysis, showing the tested H2O2 titration ranges with 0.5 mM BP in WT HEK293 cells expressing IM-APEX2, demonstrate the optimal concentration of 0.5 mM H2O2 for biotinylation experiments.

(C) WB analysis showing the distinct biotinylation pattern of HEK293 cells expressing various MICOS-APEX2 proteins and matrix-APEX2 control upon employing optimal BP (0.5 mM) and H2O2 (0.5 mM) concentration suitable for proximity labeling.

#### Supplementary Figure 2

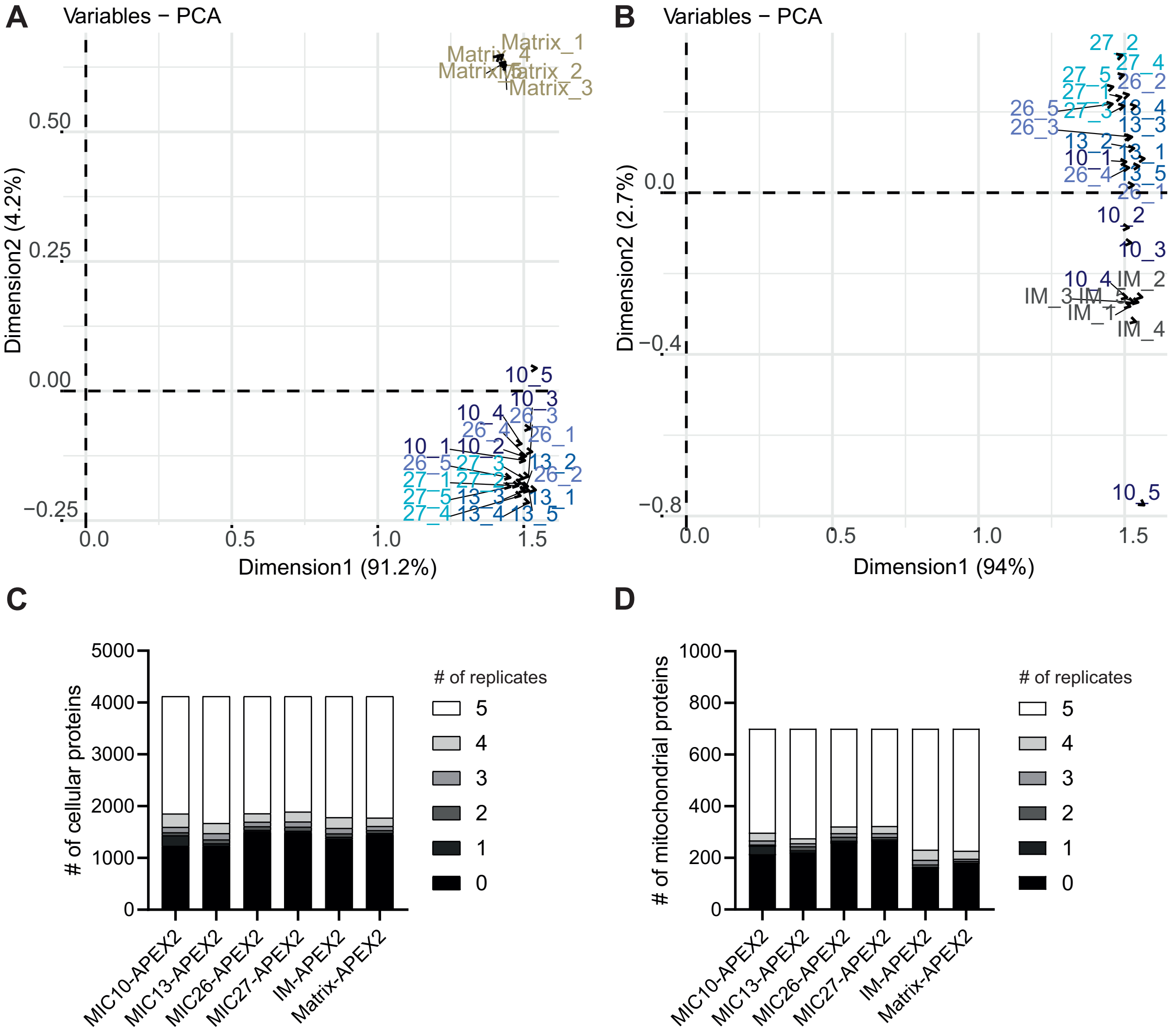


**Figure S2:** Principal component analysis plots and overview of proteins present in the number of biological replicates in mass spectrometry analysis

**Figure S2:** Principal component analysis plots and overview of proteins present in the number of biological replicates in mass spectrometry analysis

(A and B) Principal component analysis (PCA) plot, depicting the clustering of individual samples subjected to mass spectrometry, display a very close correlation between the MICOS-APEX2 fusion proteins. The matrix-APEX2 (A) manifest as distinct clusters from all the four MICOS-APEX2 fusion proteins, whereas the IM-APEX2 (B) group shows a similar distribution as the MICOS-APEX2 groups.

(C and D) Representation of the respective number of proteins detected in the corresponding number of technical replicates in MICOS-APEX2, IM-APEX2 and matrix-APEX2 controls covering the whole cell (C) and mitochondria (D).

### Supplementary Figure 3

**A**

4245 proteins detected → 4119 proteins

62 reverse
64 potential contominants

computational analysis

| **Step 1** | Cell line | MIC10-APEX2 | MIC13-APEX2 | MIC26-APEX2 | MIC27-APEX2 |
|---|---|---|---|---|---|
| detected | | 2892 | 2892 | 2613 | 2629 |
| enriched over both controls $Log_2FC$ + FOM | | 1181 | 1606 | 1146 | 1250 |
| enriched over matrix control | $Log_2FC \geq 0$ | 627 | 892 | 551 | 643 |
| | FOM | 486 | 490 | 324 | 349 |
| enriched over IM control | $Log_2FC \geq 0$ | 223 | 711 | 437 | 564 |
| | FOM | 303 | 339 | 218 | 240 |

MitoCarta3.0 filtering

| **Step 2** | Cell line | MIC10-APEX2 | MIC13-APEX2 | MIC26-APEX2 | MIC27-APEX2 |
|---|---|---|---|---|---|
| enriched over both controls $Log_2FC$ + FOM | | 206 | 239 | 176 | 178 |
| enriched over matrix control | $Log_2FC \geq 0$ | 81 | 83 | 65 | 62 |
| | FOM | 113 | 117 | 98 | 95 |
| enriched over IM control | $Log_2FC \geq 0$ | 22 | 50 | 18 | 27 |
| | FOM | 18 | 18 | 9 | 7 |

After $Log_2FC$ thresholding ($Log_2FC \geq 1$ or $Log_2FC \geq 0.5$ for matrix and IM control respectively)

| **Step 3** | Cell line | MIC10-APEX2 | MIC13-APEX2 | MIC26-APEX2 | MIC27-APEX2 |
|---|---|---|---|---|---|
| enriched over both controls $Log_2FC$ + FOM | | 168 | 192 | 152 | 151 |
| enriched over matrix control | $Log_2FC \geq 1$ | 45 | 44 | 41 | 36 |
| | FOM | 113 | 117 | 98 | 95 |
| enriched over IM control | $Log_2FC \geq 0.5$ | 10 | 28 | 13 | 20 |
| | FOM | 18 | 18 | 9 | 7 |

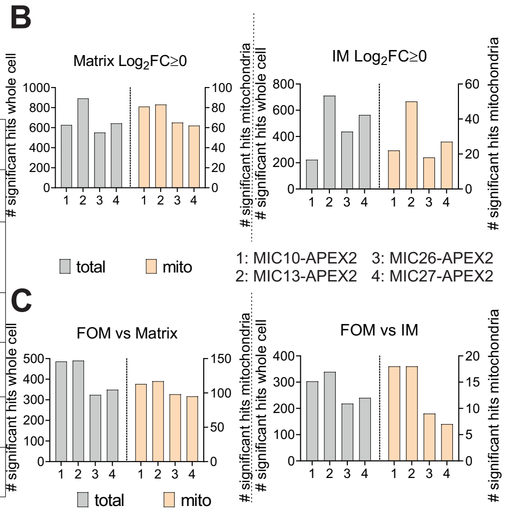


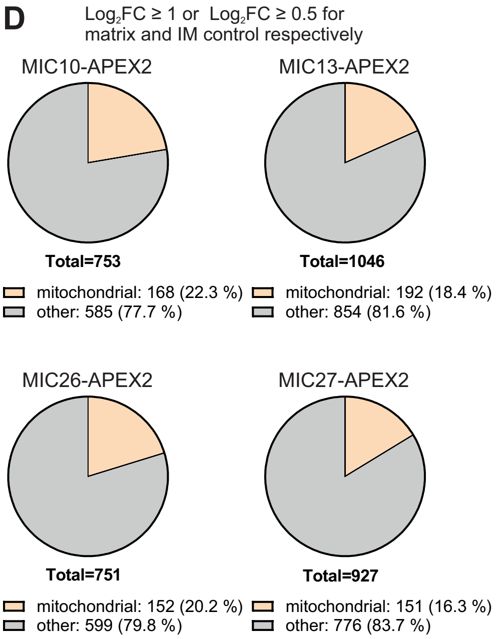


**Figure S3:** Flow chart and analysis pipeline of the interactome of all the four MICOS-APEX2 fusion proteins

**Figure S3:** Flow chart and analysis pipeline of the interactome of all the four MICOS-APEX2 fusion proteins

(A – C) Flow-chart of the analysis pipeline consisting of multiple filtering steps. The initial data set, containing 4,245 proteins, was reduced to 4,119 proteins after removal of reverse and potential contaminants. A maximum of 2613 to 2892 proteins were detected in all the four MICOS interactomes. Overall, 1146 to 1606 proteins were enriched including the log2FC enriched (increase ≥ 0) and FOM category (normalized to both the controls). The individual distribution of enriched proteins into different categories is shown (Fig 3A Step1, Fig 3B and C histograms). Post MitoCarta3.0 filtering, the number of proteins for various MICOS interactomes is shown (Step 2). Further, the enriched proteins after using log2FC cut-off of ≥ 1 for matrix-APEX2 and log2FC ≥ 0.5 for IM-APEX2 are shown (Step 3). Note the number of enriched proteins belonging to the FOM category (where no proteins were found in respective controls) does not change in step 3.

(D) Pie charts depicting the fractions of mitochondrial and non-mitochondrial proteins present in the interactome of respective MICOS proteins after the application of log2FC cut-offs of ≥1 or ≥ 0.5 for matrix-APEX2 and IM-APEX2 controls, respectively, and including the FOM candidates.

### Supplementary Figure 4

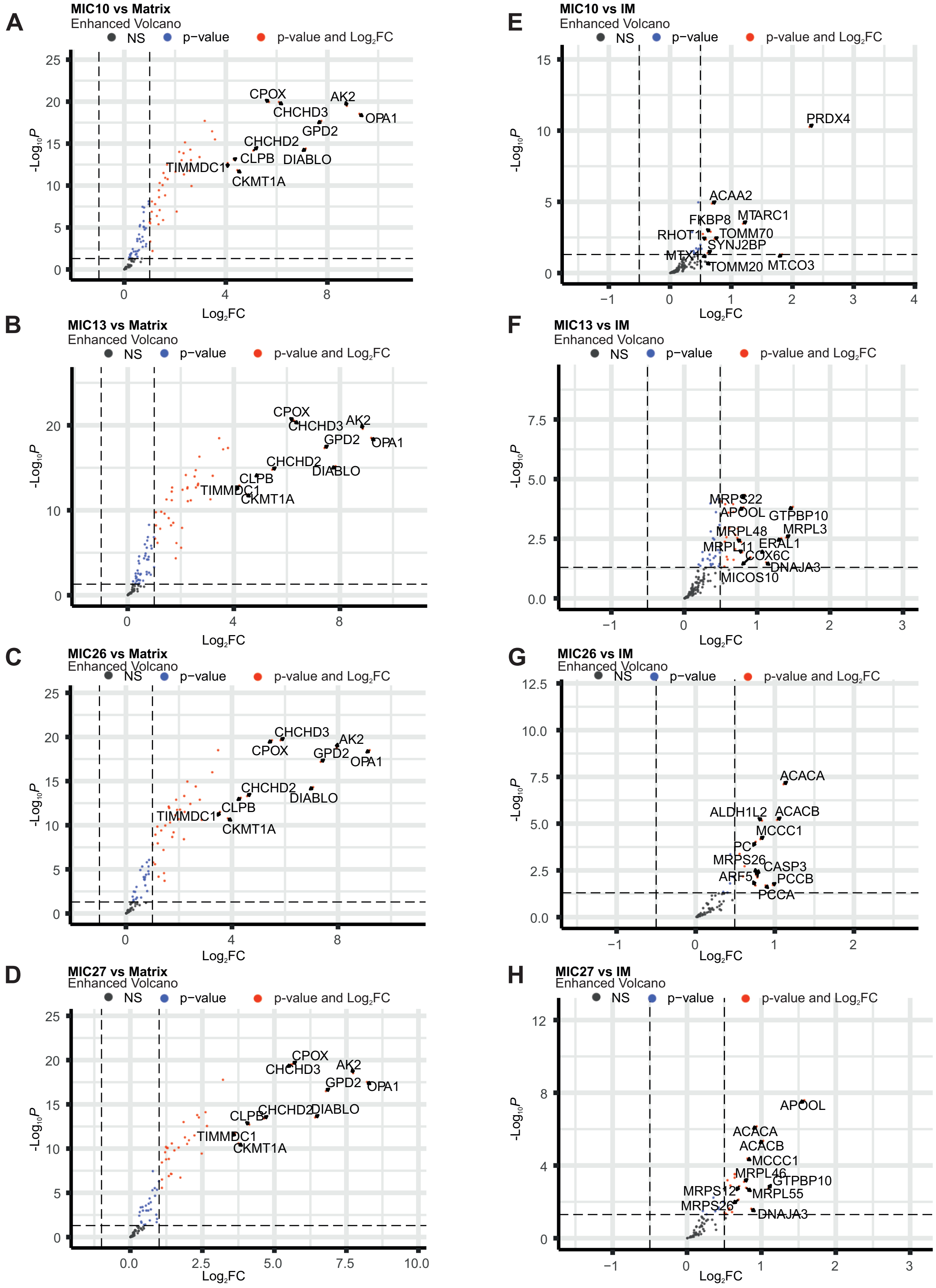


**Figure S4:** Volcano plots showing the MICOS interactors ($Log_2FC$ enriched category)

**Figure S4:** Volcano plots showing the MICOS interactors ($Log_2$FC enriched category)

(A – H) Volcano Plots showing the enriched MIC10-APEX2 (A), MIC13-APEX2 (B), MIC26-APEX2 (C), MIC27-APEX2 (D) proximity proteome when normalized to the matrix-APEX2 control as well as the IM-APEX2 control (E, F, G and H) after MitoCarta3.0 filtering. The top 10 hits with the highest log2FC enrichment are shown. OPA1 was the most enriched in all the four interactomes when compared to matrix-APEX2 control.

### Supplementary figure 5

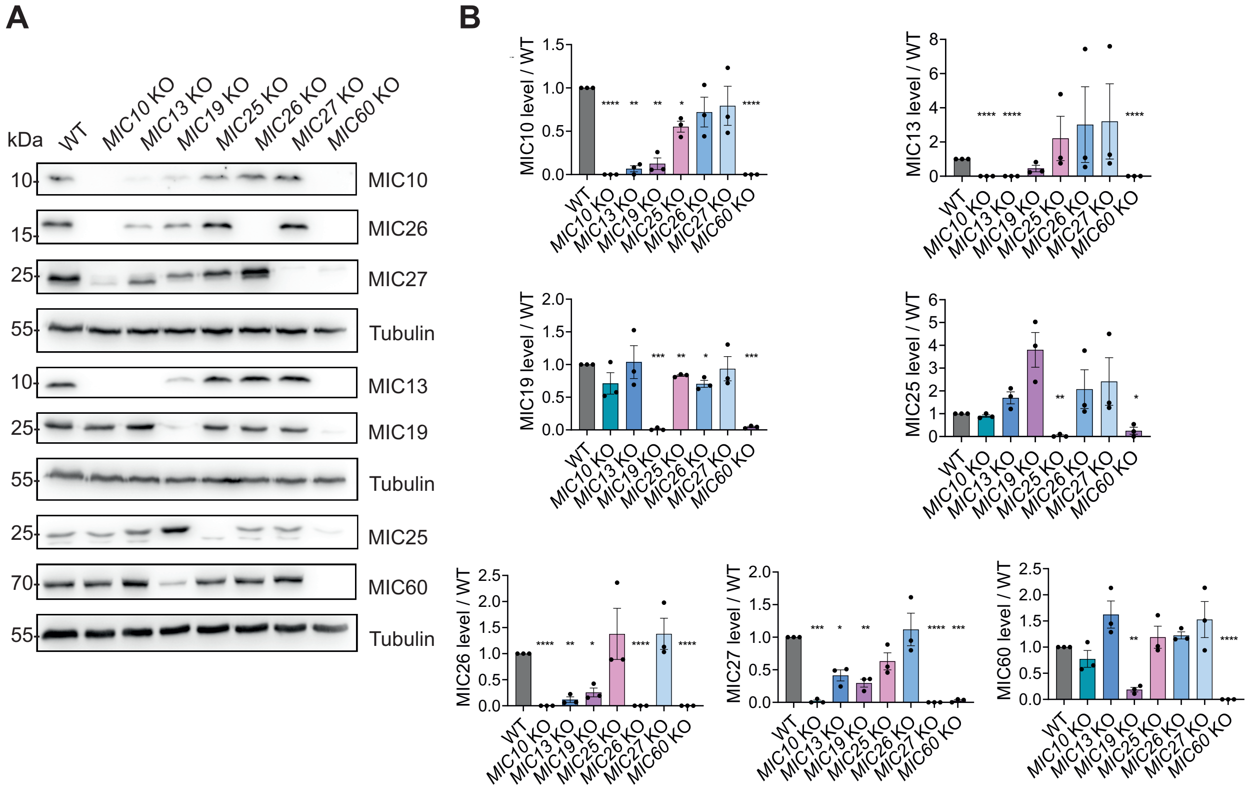


**Figure S5:** Expression of various proteins in individual MICOS HEK293 KO cells

**Figure S5:** Expression of various proteins in individual MICOS HEK293 KO cells

(A and B) Representative WB analyses of various MICOS subunits in the individual MICOS KO cell lines (A), along with the quantification from various biological replicates, reveals that deletion of either MIC10, MIC13, MIC19 or MIC60 results in total loss or strong reduction of all other MICOS subunits. Deletion of MIC25, MIC26 or MIC27 has no effect on other MICOS subunits.

Data are represented as mean ± SEM. Statistical analysis was performed using one sample t-test with *P < 0.05, **P < 0.01, ***P < 0.001, ****P < 0.0001.

#### Supplementary Figure 6

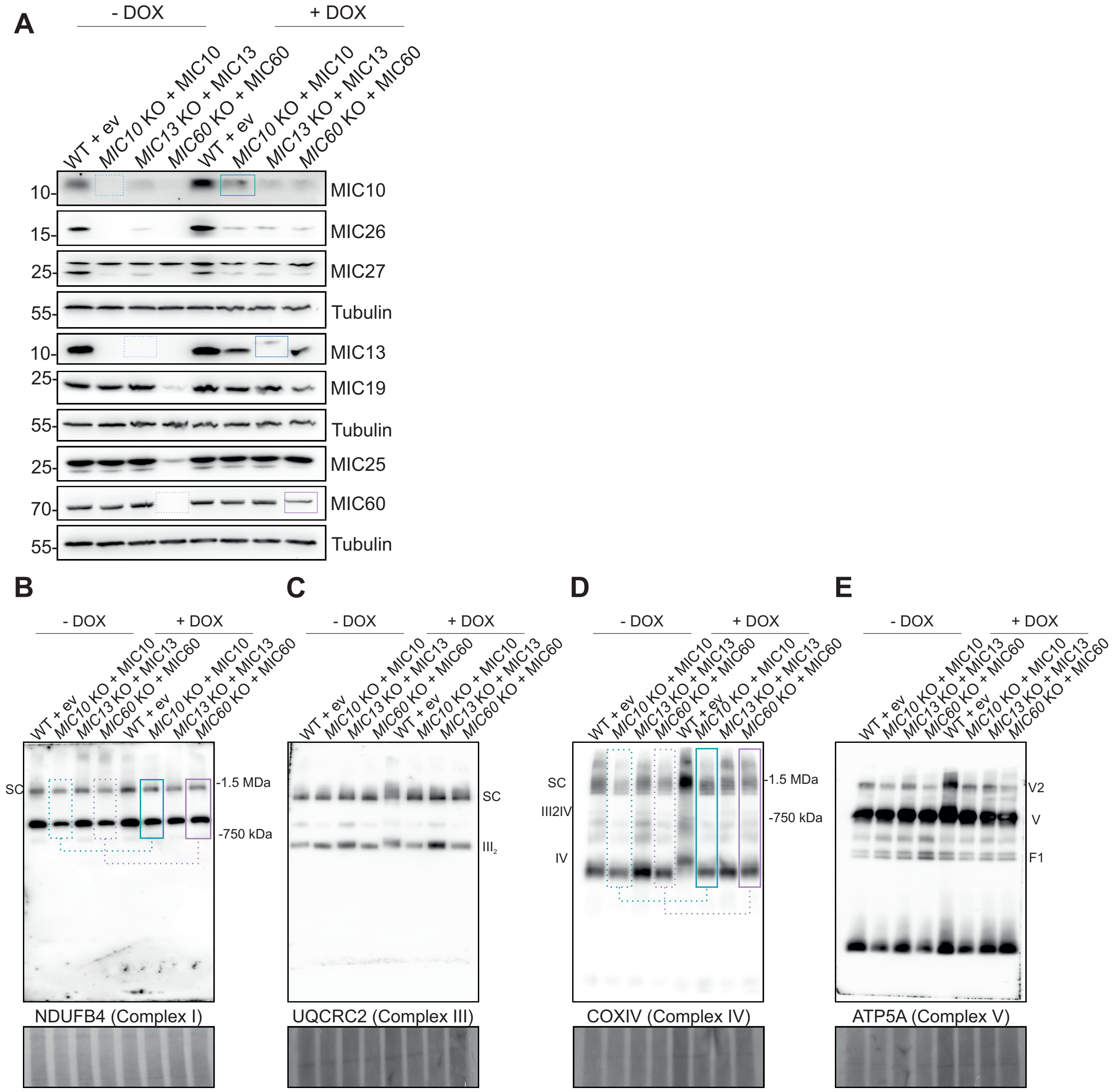


**Figure S6:** Demonstration of specificity of MIC10 and MIC60 in regulating the assembly of OXPHOS complexes I and IV

**Figure S6:** Demonstration of specificity of MIC10 and MIC60 in regulating the assembly of OXPHOS complexes I and IV

(A) WB analysis, showing the stable expression of either pLIX403-MIC10, pLIX403-MIC60 or pLIX403-MIC13-Flag along with pLIX403 empty vector control, after addition of doxycycline using the Tet-On system. Absence of MICOS proteins in corresponding MICOS KO cell lines is shown by dotted rectangles while the recovery is show using solid rectangles after doxycycline addition.

(B-E) BN-PAGE analyses of various OXPHOS complexes before and after doxycycline addition in the mentioned MICOS KO cell lines. A specific increase of OXPHOS complex I (B) and IV assembly (D) were observed after MIC10 and MIC60 induction in the Tet-On system. The rescue of OXPHOS complex I and IV assembly is shown using solid rectangles in MIC10 and MIC60 KO cells upon comparison before doxycycline addition (dotted rectangles).
